# Mutations in the *SPTLC1* gene are a cause of juvenile amyotrophic lateral sclerosis that may be amenable to serine supplementation

**DOI:** 10.1101/770339

**Authors:** J. O. Johnson, R. Chia, D. E. Miller, R. Li, Y. Abramzon, R. Kumaran, N. Alahmady, F. Faghri, A. E. Renton, S. D. Topp, H. A. Pliner, J. R. Gibbs, J. Ding, N. Smith, N. Landeck, M. A. Nalls, M. R. Cookson, O. Pletnikova, J. Troncoso, S. W. Scholz, M. S. Sabir, S. Ahmed, C. L. Dalgard, C. Troakes, A. R. Jones, A. Shatunov, A. Iacoangeli, A. Al Khleifat, N. Ticozzi, V. Silani, C. Gellera, I. P. Blair, C. Dobson-Stone, J. B. Kwok, B. K. England, E. S. Bonkowski, The International ALS Genomics Consortium, The ITALSGEN Consortium, The FALS Sequencing Consortium, The American Genome Center, P. J. Tienari, D. J. Stone, K. E. Morrison, P. J. Shaw, A. Al-Chalabi, R. H. Brown, M. Brunetti, A. Calvo, G. Mora, H. Al-Saif, M. Gotkine, F. Leigh, I. J. Chang, S. J. Perlman, I. Glass, C. E. Shaw, J. E. Landers, A. Chiò, T. O. Crawford, B. N. Smith, B. J. Traynor

## Abstract

Juvenile amyotrophic lateral sclerosis (ALS) is a rare form of childhood motor disorder with a heterogeneous clinical presentation. The underlying causes of this condition are poorly understood, hindering the development of effective therapies. In a whole-exome sequencing trio-family study of three unrelated juvenile patients diagnosed with ALS and failure to thrive, we identified de-novo mutations in *SPTLC1* (p.Ala20Ser in two patients and p.Ser331Tyr) not present in their healthy parents or siblings. *SPTLC1* encodes a subunit of the serine palmitoyltransferase complex, a key enzyme in sphingolipid biosynthesis. Mutations in this gene are known to cause hereditary sensory autonomic neuropathy, type 1A, with a characteristic increase in plasma levels of neurotoxic deoxymethyl-sphinganine. We found an increase of this metabolite in one of our patients carrying the p.Ala20Ser mutation. Treatment of one of the patients with high dose, oral L-serine led to an increase in body weight, suggesting that serine supplementation may be beneficial among patients carrying mutations in this gene.

## Introduction

Amyotrophic lateral sclerosis (ALS, OMIM number 105400) is a relatively common neurological disorder characterized by rapidly progressive paralysis leading to death from respiratory failure. The mean age of onset of ALS is the seventh decade of life, with the vast majority of cases occurring after the age of forty (*1*). In contrast, juvenile ALS (defined as an age of onset less than twenty-five years of age) is a rare form of motor neuron disease (*2*). These early-onset cases are characterized by relatively slow progression and a variable phenotype that often makes accurate diagnosis challenging (*2*).

Considerable progress has been made in unravelling the genetic architecture underlying ALS, but much remains to be understood about this condition (*3*). For example, genome-wide association studies have identified only a handful of loci, many of which have proven difficult to replicate (*3*). Our lack of knowledge concerning the pathogenesis of ALS hampers efforts to design disease-modifying interventions. Juvenile ALS is thought to be more frequently genetic in origin than the adult-onset forms, and the genetic analysis of these young-onset cases offers an opportunity to identify disease-causing genes (*2*). By extension, any gene underlying juvenile ALS may also play a role in adult-onset cases.

De-novo genetic variants may underlie at least a portion of ALS cases. Such mutations would not be readily detected by genome-wide association studies owing to their recent occurrence and corresponding low frequency within the community. Spontaneously occurring mutations are a well-known cause of neurological conditions, such as neurofibromatosis type 1 and Duchenne muscular dystrophy (*4*, *5*). Indeed, de-novo mutations of the familial ALS genes, *FUS*, *SOD1*, and *VCP*, have been previously described in sporadic ALS cases (*6*, *7*). Such spontaneous mutations are more likely to present with early-onset disorders due to their impact on fitness (*8*).

Here, we performed whole-exome sequencing of three patients diagnosed with juvenile ALS and their unaffected parents, to identify the mutations responsible for their disease. None of these cases had a family history of ALS or neuromuscular disorders, suggesting de-novo variations as the underlying genetic mechanism. We also provide preliminary evidence that serine supplementation could revert features of mitochondrial abnormalities in heterologous cell models expressing mutant SPTLC1. Importantly, we demonstrate that oral L-serine significantly improved one of our patient’s failure to thrive, which is a promising early clinical proxy following the natural course of the disease.

## Results

### Identification of *de novo* mutations in *SPTLC1* as a cause of juvenile ALS

We performed whole-exome sequencing of three unrelated patients and their healthy parents who had been diagnosed with juvenile ALS (**Figure 1, Table 1**). The severe phenotype observed among these patients, together with the lack of a family history of neurological diseases (**Figure 2A-C**), suggested spontaneous mutations as the underlying genetic defect. The analysis of their genetic data identified de-novo variants in the serine palmitoyltransferase long chain base subunit 1 (*SPTLC1*) gene that were present in each of the three patients and were absent in their parents (**Figure 2D**). Patients 1 and 2 carried the same heterozygous p.Ala20Ser mutation in *SPTLC1* due to variation in adjacent nucleotides (chr9:94874844, C>A and chr9:94874843, G>T). Patient 3 carried a p.Ser331Tyr (chr9:92047261G>T) heterozygous mutation in *SPTLC1*. These *SPTLC1* mutations were not present in control subjects or online databases of human polymorphisms (n = 142,489 individuals). The p.Ser331Tyr amino acid change has been previously implicated in neurological disease (*9*), whereas the p.Ala20Ser mutations have not been reported previously.

**Figure 1.**
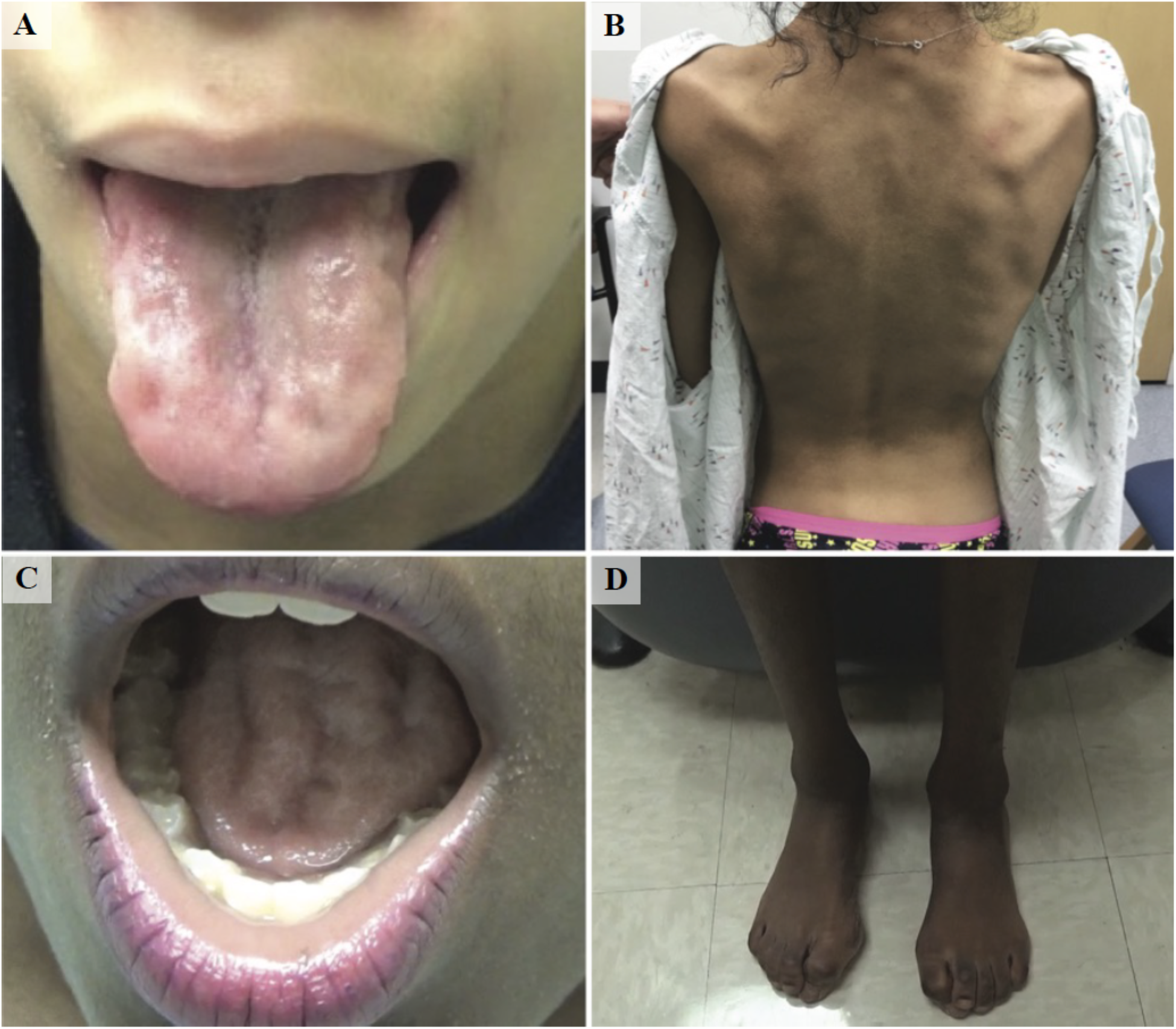
Clinical features of the patients diagnosed with juvenile ALS. (**A & B**) Tongue wasting and scapular winging in patient 2 carrying the p.Ala20Ser *SPTLC1* mutation. (**C &D**) Tongue wasting and muscle atrophy of the lower limbs in patient 3 carrying the p.Ser331Tyr *SPTLC1* mutation. Note the hammertoe deformities of both feet.

**Table 1.** Clinical features of patients diagnosed with juvenile ALS and carrying de-novo mutations in *SPTLC1*.

| Clinical features | Patient 1 | Patient 2 | Patient 3 |
| --- | --- | --- | --- |
| Gene change | p.Ala20Ser | p.Ala20Ser* | p.Ser331Tyr |
| Age at onset | 5 yrs | 8 yrs | 4 yrs |
| Age at evaluation | 20 yrs | 15 yrs | 11 yrs |
| BMI (Z-score) | 13th pctl (-1.12) | <1<sup>st</sup> pctl (-7.0) | <1<sup>st</sup> pctl (-6.5) |
| Back deformities | Severe scoliosis | Lordosis | Normal posture |
| Foot deformities | Pes cavus | - | Pes cavus/varus |
| Walking | Non-ambulatory | Steppage | Steppage |
| Atrophy | Global, contractures | Global | Global |
| Weakness | Generalized | Generalized | Generalized |
| Reflexes | Hyporeflexia,<br>Ach brisk | Hyporeflexia,<br>Ach brisk | Hyperreflexia,<br>Ach absent |
| Tongue | Wasted and<br>fasciculating | Wasted and<br>fasciculating | Wasted and<br>fasciculating |
| Jaw jerk | Present | - | Present |
| Respiratory | Trach. at 17 yrs | - | Dyspnea on exercise |
| Cognition | Frontal lobe<br>dysfunction | Executive dysfunction | Normal |
| Sensory | Normal | Normal | Moderate pain loss in<br>glove-stocking, painless<br>foot ulceration |
| Neurophysiology |  |  |  |
| Motor | Chronic denervation | Acute and chronic<br>denervation | Axonal loss<br>Polyphasia |
| Sensory | Normal | Normal | Axonal loss |
| Additional features | - | Scapular winging,<br>Gower's sign | Gower's sign, vitamin D<br>def., hyperhidrosis |

\*variant was detected in 1 of 49 and 0 of 149 next-generation sequencing reads from the father's saliva-derived and
buccal-derived DNA. Ach, Achilles tendon; yrs, years; trach, tracheostomy; def., deficiency; pctl, percentile.

**Figure 2.**
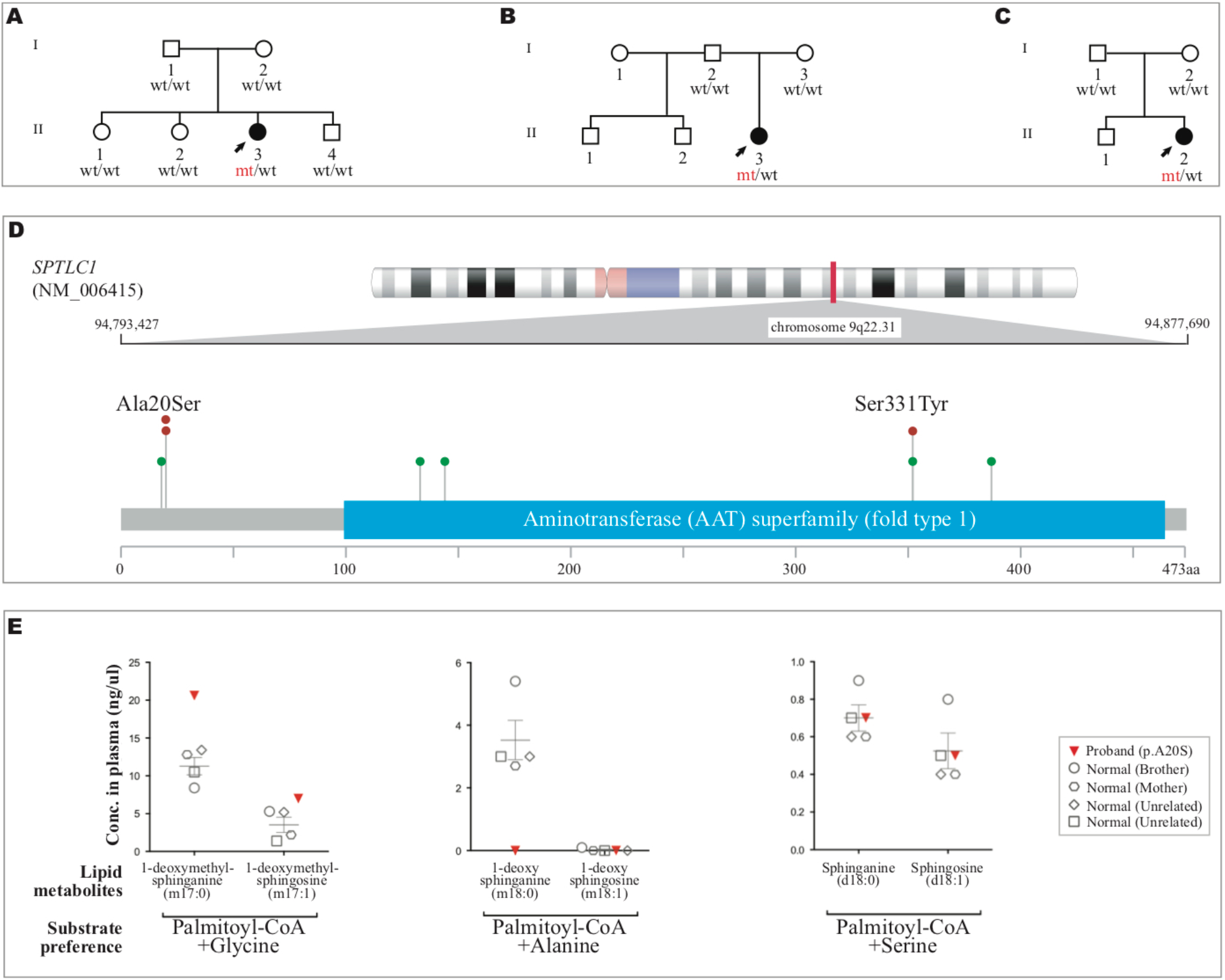
De-novo mutations of *SPTLC1* in patients diagnosed with juvenile ALS. **(A-C)** Pedigree of three patients diagnosed with juvenile ALS. The mutant alleles in *SPTLC1* are indicated by *mt*, whereas wild-type alleles are indicated by *wt*. The arrows indicate the probands. **(D)** Distribution of *SPTLC1* mutations detected in patients diagnosed with juvenile ALS. Mutations identified in the three juvenile ALS cases are noted in red, and mutations previously described to cause HSAN1 are shown in green. **(E)** Plasma analysis of the proband and healthy controls (unaffected mother, unaffected brother, and two unrelated healthy individuals). Proband had ~1·8 times higher levels of 1-deoxymethyl-sphinganine and 1-deoxymethyl-sphingosine compared to the healthy controls. The utilization of L-alanine by the serine palmitoyltransferase complex produces 1-deoxysphinganine (m18:0) and 1-deoxysphingosine (m18:1), whereas the use of L-glycine yields 1-deoxymethyl-sphinganine (m17:0) and 1-deoxymethyl-sphingosine (m17:1). Collectively, these non-canonical metabolites are called deoxysphingoid bases.

### Elevated plasma levels of deoxymethyl-sphinganine in a patient carrying p.Ala20Ser *SPTLC1* mutation

Mutations in *SPTLC1* are a known cause of autosomal dominant hereditary sensory and autonomic neuropathy, type 1A (HSAN1, OMIM number 162400) (*10*, *11*). The protein encoded by *SPTLC1* is an essential subunit of serine palmitoyltransferase (SPT), the enzyme that catalyzes the first and rate-limiting step in the de-novo synthesis of sphingolipids (*12*). A characteristic feature of *SPTLC1* mutations associated with HSAN1 is a shift in substrate specificity of serine palmitoyltransferase to L-alanine and L-glycine, leading to the formation of an atypical class of deoxy-sphingolipids (*13*). These neurotoxic metabolites accumulate within cells as they cannot be converted to complex sphingolipids nor degraded by the catabolic pathway (*13*). Abnormally elevated plasma levels of deoxy-sphingolipids are a known biomarker for pathogenic *SPTLC1* mutations in HSAN1 patients (*13*).

Based on this information, we used mass spectrometry to test for this biomarker in patient 1 carrying the p.Ala20Ser *SPTLC1* mutation, and found an abnormally high level of 1-deoxymethyl-sphinganine in her plasma that was not present in her unaffected parents, her unaffected siblings, or control subjects (**Figure 2E** and **Table S2**). These findings confirmed that this de-novo mutation had abnormally altered the function of the SPTLC1 protein and was the likely cause of disease in this individual.

### Serine supplementation ameliorates damaging effects caused by mutant p.Ala20Ser

Using a photometric assay of SPT enzyme activity, we found that the p.Ala20Ser mutant SPTLC1 complex has an altered preference for L-alanine and glycine over the canonical L-serine, compared to the wild-type SPTLC1 complex (**Figure S1**). Similar altered kinetics were also observed from cellular-based assays using known HSAN1-mitochondrial phenotypes (membrane polarization and size) as readouts (*14*). Measurements were benchmarked against defects found in cells expressing the known HSAN1 mutation p.Cys133Trp that served as an internal positive control. Mitochondrial size and intensity were defective to the same degree in cells expressing p.Ala20Ser and p.Cys133Trp and was reversed to the wild-type phenotype upon the supplementation of serine in culture (**Figure S1**).

Based on our findings and work published by other groups(*13*, *15*), and in light of the poor prognosis observed among juvenile ALS patients, patient 2 was commenced on high-dose oral serine supplementation. She began taking 6 grams per day for five and a half weeks, followed by 9 grams per day for two and a half weeks, and presently is ingesting 10 grams per day. Treatment was well tolerated over six months, and her body weight increased after having stagnated for years (**Figure 3**). Thus far, we did not observe objective evidence of neurological improvement.

**Figure 3.**
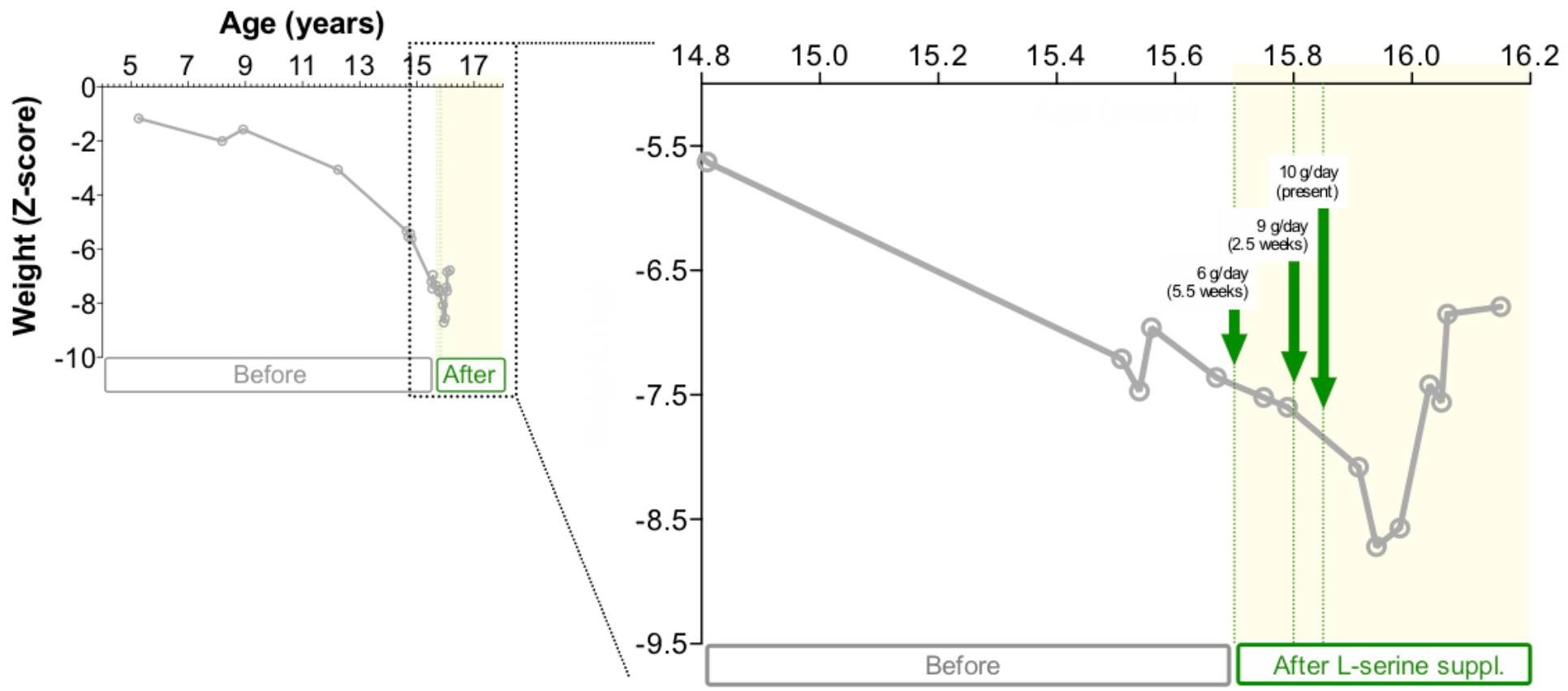
Body mass index progression of patient 2 on high-dose oral serine supplementation. The weight profile of patient 2 indicated a steady loss in weight until she was started on high-dose oral serine supplementation at the age of 15.7 years. Weights are presented as Body mass index (BMI) z-scores that are measures of relative weight adjusted for child age and sex. Higher values represent higher BMI. The vertical dashed lines indicate dose increases. Treatment was well tolerated over six months, and her weight increased, surpassing her maximum recorded weight.

### *SPTLC1* variants in adult-onset ALS patients are rare

Having established that mutations in *SPTLC1* are a cause of juvenile ALS, we explored the role of mutations in this gene in the pathogenesis of adult-onset ALS. To do this, we evaluated the occurrence of *SPTLC1* mutations in a series of 5,607 cases with adult-onset ALS. This screening identified 17 novel *SPTLC1* mutations in 19 (0·34%) ALS cases that were rare or absent in healthy subjects and were predicted to be damaging (**Table S3, Figure S2** and **S3**). Gene burden testing showed a trend towards significance (87 variants in population samples, one-sided Fisher test p-value using TRAPD software package = 0.0014). Due to the late age at presentation of the adult ALS patients, DNA was not available from their parents to determine whether these variants occurred spontaneously. The typical clinical features of ALS were observed among these adult-onset patients, though this subgroup was more likely to report a family history of ALS (n = 5 patients, 26·3%, **Table 2**). None of the patients reported sensory or autonomic involvement. The intensity and number of motor neurons staining with SPTLC1 were diminished in autopsy tissue obtained from an ALS patient carrying a p.Arg445Gln mutation in *SPTLC1* (**Figure S4**).

**Table 2.**
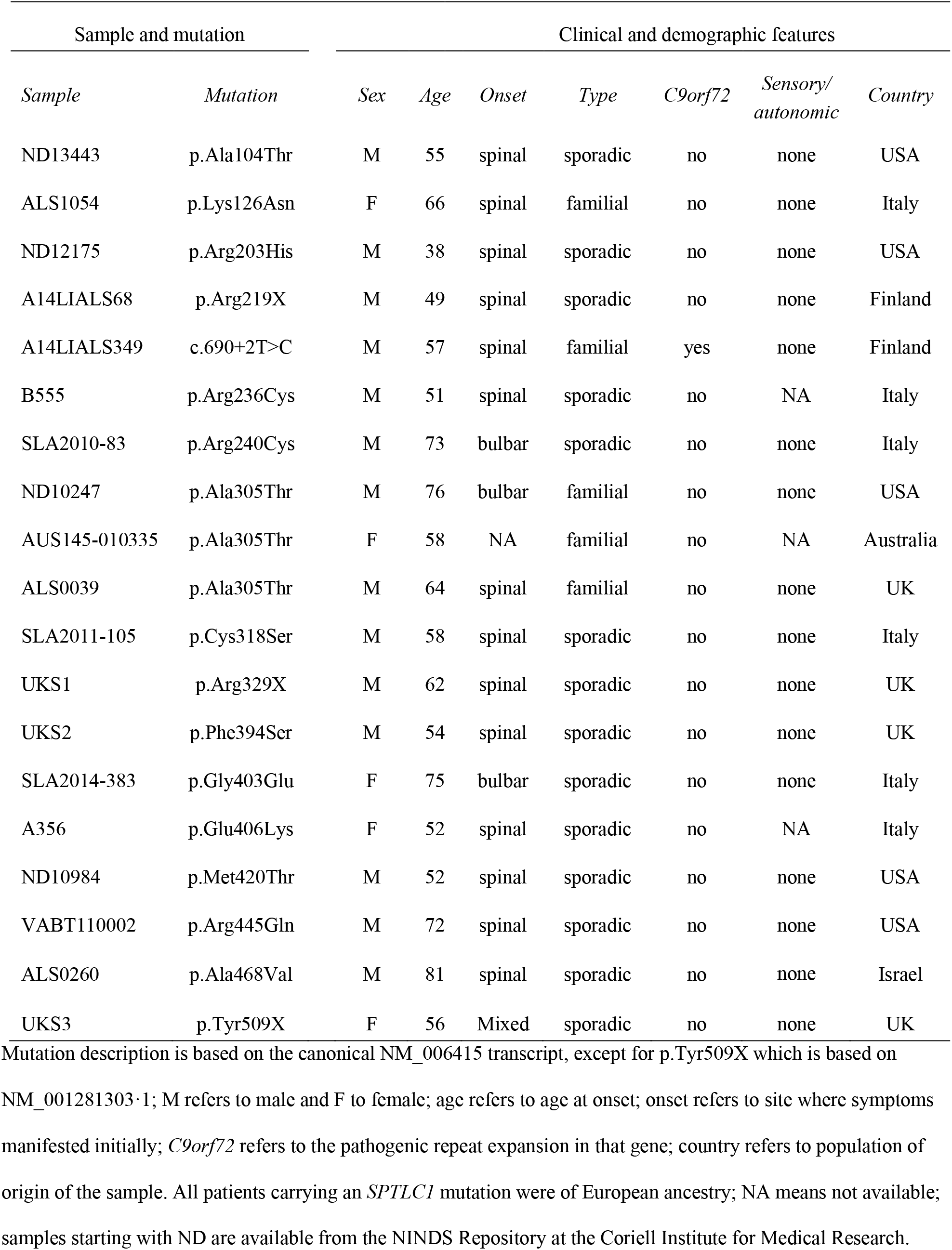
Demographic and clinical features of patients diagnosed with ALS carrying *SPTLC1* mutations.

| Sample and mutation |  | Clinical and demographic features |  |  |  |  |  |  |
| --- | --- | --- | --- | --- | --- | --- | --- | --- |
| *Sample* | *Mutation* | *Sex* | *Age* | *Onset* | *Type* | *C9orf72* | *Sensory/*<br>*autonomic* | *Country* |
| ND13443 | p.Ala104Thr | M | 55 | spinal | sporadic | no | none | USA |
| ALS1054 | p.Lys126Asn | F | 66 | spinal | familial | no | none | Italy |
| ND12175 | p.Arg203His | M | 38 | spinal | sporadic | no | none | USA |
| A14LIALS68 | p.Arg219X | M | 49 | spinal | sporadic | no | none | Finland |
| A14LIALS349 | c.690+2T>C | M | 57 | spinal | familial | yes | none | Finland |
| B555 | p.Arg236Cys | M | 51 | spinal | sporadic | no | NA | Italy |
| SLA2010-83 | p.Arg240Cys | M | 73 | bulbar | sporadic | no | none | Italy |
| ND10247 | p.Ala305Thr | M | 76 | bulbar | familial | no | none | USA |
| AUS145-010335 | p.Ala305Thr | F | 58 | NA | familial | no | NA | Australia |
| ALS0039 | p.Ala305Thr | M | 64 | spinal | familial | no | none | UK |
| SLA2011-105 | p.Cys318Ser | M | 58 | spinal | sporadic | no | none | Italy |
| UKS1 | p.Arg329X | M | 62 | spinal | sporadic | no | none | UK |
| UKS2 | p.Phe394Ser | M | 54 | spinal | sporadic | no | none | UK |
| SLA2014-383 | p.Gly403Glu | F | 75 | bulbar | sporadic | no | none | Italy |
| A356 | p.Glu406Lys | F | 52 | spinal | sporadic | no | NA | Italy |
| ND10984 | p.Met420Thr | M | 52 | spinal | sporadic | no | none | USA |
| VABT110002 | p.Arg445Gln | M | 72 | spinal | sporadic | no | none | USA |
| ALS0260 | p.Ala468Val | M | 81 | spinal | sporadic | no | none | Israel |
| UKS3 | p.Tyr509X | F | 56 | Mixed | sporadic | no | none | UK |

Mutation description is based on the canonical NM\_006415 transcript, except for p.Tyr509X which is based on
NM\_001281303.1; M refers to male and F to female; age refers to age at onset; onset refers to site where symptoms manifested initially; *C9orf72* refers to the pathogenic repeat expansion in that gene; country refers to population of origin of the sample. All patients carrying an *SPTLC1* mutation were of European ancestry; NA means not available; samples starting with ND are available from the NINDS Repository at the Coriell Institute for Medical Research.

Among the adult-onset ALS cases, we found the p.Ala305Thr variant in *SPTLC1* in three patients with familial ALS. These individuals shared a common haplotype extending across the locus, suggesting that they share a common ancestor (**Table S5**). Though rare (n = 2 out of 135,025 population carried the same SNP), this variant did not segregate within a large, Australian ALS-FTD kindred (**Figure S5**) (*16*). Furthermore, a missense mutation in the gene *CYLD* has recently been nominated as the cause of disease in this family (*17*). Additionally, two siblings in a UK kindred carried the p.Ala305Thr variant and remained healthy in their seventh decade of life. Overall, these observations point toward p.Ala305Thr being either a non-pathogenic variant or a rare, variably penetrant risk factor.

## Discussion

We provide compelling genetic, biochemical, and cellular data that mutations in *SPTLC1* are a cause of juvenile ALS. First, we found three unrelated patients diagnosed with juvenile ALS who carried de-novo mutations in this gene. These variants were not present in our in-house control dataset or online databases of human polymorphisms, indicating they were rare variants in ethnically diverse populations. Two of the patients carried the same alanine to serine amino acid shift at position 20 of the protein which were from different nucleotide changes. Second, mass spectrometry analysis of plasma showed the presence of neurotoxic deoxy-sphingolipid metabolites confirming that the observed mutation alters the function of the encoded enzyme, leading to increased aberrant utilization of alanine and glycine as substrates. This biochemical pattern was consistent with a previously reported disease-causing mechanism in HSAN1 patients (*13*). Third, we used immunohistochemistry to demonstrate that SPTLC1 is abundantly expressed within the motor neurons of healthy spinal cord tissue.

Though labeled as hereditary sensory and autonomic neuropathy type 1A, the phenotypes associated with mutations in *SPTLC1* are known to be varied with patients manifesting various combinations of sensory loss, autonomic dysfunction, and motor weakness (*18*). Indeed, there is a previous report of a de-novo p.Ser331Phe mutation in *SPTLC1* in a young French girl presenting with a similar phenotype to our patients. Her clinical picture consisted of severe restriction of growth, cognitive impairment, amyotrophy, hyperreflexia, vocal cord paralysis, and respiratory failure, though this patient was not formally diagnosed as having juvenile ALS (*9*). More recently, retinal disease has been reported in patients carrying *SPTLC1* mutations (*19*). This clinical heterogeneity has been linked to the differing effects of each mutation on SPTLC1 enzyme-substrate preference (*13*), and we observed similar differences in substrate utilization across the variants that we had studied at the enzymatic level (**Figure S1**).

Perturbed sphingolipid metabolism underlies many neurological disorders, such as Niemann-Pick disease and Gaucher disease (*20*), and may play a role in the pathogenesis of Alzheimer’s disease (*21*). Sphingolipid metabolism has also been directly implicated in motor neuron degeneration. For example, patients with partial deficiency of hexosaminidase A enzyme activity (also known as GM2 gangliosidosis, a form of sphingolipidosis) may have clinical manifestations mimicking ALS (*22*). Accumulation of ceramides and cholesterol esters also occurs within the spinal cords of ALS patients and a *SOD1* transgenic mouse model of ALS (*23*).

Our data suggest that patients diagnosed with juvenile ALS should be screened for *SPTLC1* mutations, especially as dietary supplementation with serine may be beneficial if instituted at an early stage in the disease course (*15*). Indeed, the treatment of one of our patients with serine resulted in significant weight gain, which was the first time she had gained weight in several years and an initial proof of concept. We did not observe evidence of neurological improvement, though we recognize that prolonged therapy would be required to detect such an effect (*24*). Serine is a non-essential amino acid that is widely available as a low-cost nutritional supplement. A 10% serine-enriched diet reduced neurotoxic deoxysphingolipid plasma levels both in transgenic mice expressing the p.Cys133Trp *SPTLC1* mutation and in human patients diagnosed with HSAN1 (*15*). Furthermore, a safety trial involving twenty adult-onset ALS patients demonstrated that high doses of oral serine are well-tolerated and that this polar amino acid is actively transported across the blood-brain barrier (*25*). Nutritional supplementation has proven to be remarkably effective in other forms of ALS: high dose oral vitamin B_2_ (riboflavin) slows and even halts neurological progression in Brown-Vialetto-Van Laere cases, a rare subtype of childhood or young-onset ALS arising from mutations in the riboflavin pathway (*26*).

Our study has limitations. Our evidence suggesting that mutations in *SPTLC1* are a cause of adult-onset ALS is preliminary. Though significance was achieved based on the testing of a single gene, our gene burden testing did not reach genome-wide significance, as the p-value for *SPTLC1* in adult-onset ALS cases corrected for multiple testing of 20,000 genes was 1.0. In part, this may be because of the rarity of these mutations and because we could not show segregation of the disease within extended kindreds of the p.Ala305Thr variant. The lack of large pedigrees, due to the poor survival associated with the disease, has hampered the ALS genetics field. Even if mutations in *SPTLC1* are confirmed to be a cause of adult-onset ALS by future studies, only a small number of patients diagnosed with ALS are likely to carry pathogenic *SPTLC1* mutations. Despite this rarity, the lack of effective treatments for ALS suggests that it may be worthwhile to screen adult ALS patients for *SPTLC1* mutations. Serine nutritional supplementation is cheap (~$500 per year) and relatively safe (*25*). In such cases, abnormal deoxysphingolipid plasma metabolites could be used as a diagnostic biomarker to confirm the toxicity of *SPTLC1* mutations, and as a marker of target engagement after commencing serine treatment (*27*).

In conclusion, our data broaden the phenotype associated with mutations in *SPTLC1* to include juvenile ALS and implicates sphingolipid metabolism as a pathway in motor neuron disease. Our findings are particularly relevant in light of the fact that nutritional supplementation with serine may ameliorate the toxic effect of abnormal sphingolipid metabolites in these patients if instituted at an early stage in the disease. This provides an early opportunity to test the precision medicine approach in an otherwise fatal neurodegenerative disease.

## Materials and Methods

### Patients

Three unrelated patients with neuromuscular symptoms consistent with juvenile ALS took part in the study between March 2016 and March 2020. **Table 1** summarizes the clinical features of the three patients. Recruitment information and a detailed description of each case with videos of pertinent neurological findings are available in the appendix.

Patient 1 presented with gradually progressive spastic diplegia and growth retardation at five years of age. By age twenty, she had quadriplegia with marked muscle atrophy and diminished weight, brisk lower limb reflexes, tongue fasciculations and weakness, dysarthria, mild cognitive dysfunction, and respiratory failure requiring tracheostomy and ventilation. Repeated neurophysiological testing did not show evidence of sensory or autonomic dysfunction. She was diagnosed with juvenile ALS based on the revised El Escorial criteria (*28*).

Patient 2 was a fifteen-year-old, right-handed girl with mixed African American and white race/ethnicity who presented with a six-year history of gradually progressive generalized limb and bulbar weakness. She had a long-standing history of diminished weight of unknown cause, and her school performance began to decline at the age of fifteen. Her neurological examination at presentation revealed a body mass index less than the 1^st^ percentile, exaggerated lumbar lordosis, tongue fasciculations and wasting, generalized muscle atrophy and weakness, brisk asymmetric ankle reflexes, a positive Gower’s sign, and normal sensation (**Figure 1A-B, Movie S1 and S2**). Neurophysiological testing revealed active and chronic denervation without evidence of sensory neuropathy. Decreased sustained attention and impaired executive functioning was evident on neuropsychological evaluation. She was diagnosed with juvenile ALS based on the revised El Escorial criteria (*28*).

Patient 3 was an eleven-year-old African American girl with a history of failure to gain weight and toe-walking since the age of four. She presented at age ten with a deteriorating gait, hand weakness, right foot paresthesia, dysphagia, and increased sweating. Examination revealed marked atrophy, postural tachycardia, bilateral cataracts, a wasted, fasciculating tongue with an exaggerated jaw jerk, generalized fasciculations and weakness associated with hyperreflexia, and decreased pinprick sensation in a glove-and-stocking distribution (**Figure 1C-D, Movie S3**). The patient walked abnormally due to weakness and bilateral foot drop, and she had a positive Gower’s sign. Neurophysiological examination showed sensorimotor axonal neuropathy, as well as polyphasic potentials on electromyography. She was diagnosed with a juvenile ALS-plus syndrome due to her prominent motor symptoms and modest sensory-autonomic involvement. The pedigrees of patients 1, 2 and 3 are shown in **Figure 2A-C**.

For mutational screening of *SPTLC1* in adult-onset ALS, we used 5,607 DNA samples that were obtained from individuals diagnosed with adult-onset ALS (**Table S1**). Control data consisted of 5,710 neurologically healthy U.S. individuals who had undergone next-generation sequencing at the Laboratory of Neurogenetics or as part of the Alzheimer’s Disease Sequencing Project (ADSP). All subjects provided written informed consent for genetic analysis according to the Declaration of Helsinki, and the Institutional Review Board of the National Institutes of Health approved the study.

### Next-generation sequencing in juvenile ALS

Whole-exome sequencing was performed using 100 base pair, paired-end sequencing on an Illumina sequencer according to the manufacturer’s protocol. DNA from patient 1 and her family was sequenced in the Laboratory of Neurogenetics using TruSeq library preparation (version 1.0). DNA from patients 2 and 3 and their families was sequenced at GeneDx using IDT xGen Exome Research Panel (version 1.0) (*29*).

Data were analyzed to identify de-novo variants present in the affected child, and not present in either parent. As the mutations underlying a rare disease, such as ALS, are unlikely to be present in the general population, variants present in the Genome Aggregation Database (gnomAD, version 2·1), or the Kaviar Genomic Variant database (version September 23, 2015) were excluded. Synonymous, intronic, and intergenic changes were similarly excluded (*ANNOVAR*, August 11, 2016 version). Paternity and maternity were confirmed using identity-by-descent analysis, and exome data were reviewed to identify mutations in known ALS genes.

### *SPTLC1* sequencing in adult-onset ALS

DNA from adult-onset ALS cases were sequenced to identify mutations in the *SPTLC1* gene. The sequence data was generated using whole-exome sequencing (n = 3,683 cases) (*30*, *31*), whole-genome sequencing (n = 1,274 cases), and Sanger sequencing (n = 650) of the fifteen exons and flanking introns of *SPTLC1* using the primer sequences listed in **Table S4**. Variants detected in *SPTLC1* based on whole-genome sequence data were confirmed by Sanger sequencing.

Variants in *SPTLC1* were considered to be deleterious if they: (a) were not present in the 4,647 control ADSP subjects; (b) had a frequency less than 3·3×10^−5^ in online databases of human polymorphisms including the 51,592 European and 8,949 Finnish non-neurological individuals in gnomAD, and the 77,301 samples in Kaviar;^1^ and (c) were designated as ‘damaging’ according to four out of five prediction algorithms (*32*); were identified as “stop gain” or “frameshift”; or as splice site mutations with a dbscSNV score higher than 0·6. Gene burden testing of *SPTLC1* was performed using publicly available control data (gnomAD and Kaviar) as implemented in the Test Rare vAriants with Public Data (TRAPD) software package (*33*). This script performs a one-sided Fisher’s exact test to determine if there is a higher burden of qualifying variants in cases as compared to controls for a tested gene. The threshold for statistical significance was set at p ≤ 0.05.

### Measurement of sphingolipid levels

Sphingolipids were extracted from plasma using methanol extraction (*34*). Chromatographic separation was performed on an Acquity Ultra Performance Liquid Chromatography system (Waters), and sphingolipids were detected in a Xevo TQ-S triple quadrupole-mass spectrometer (Waters) using the multiple reaction monitoring method. Acquired data were analyzed using *MassLynx* software (version 4·1). Calibration equations for the sphingolipids were obtained by plotting response against concentration (ng/mL). The equation showed good linearity over the 1 ng to 3,000 ng range.

## Supporting information

Movie S1. Tongue fasciculations and wasting in patient 2

Movie S3. Neurological manifestations in patient 3

Movie S4. MRI brain and spinal cord in patient 2

## General

We thank the patients and their families for their participation in our research. We also thank Kirsty McWalter and GeneDx for their assistance. This study used DNA samples, genotype data, and clinical data from the NINDS Repository at Coriell, the New York Brain Bank-The Taub Institute, Columbia University, Department of Veterans Affairs Biorepository Brain Bank (grant #BX002466), the Baltimore Longitudinal Study of Aging, the Johns Hopkins University Alzheimer’s Disease Research Center (NIH grant P50AG05146), the NICHD Brain and Tissue Bank for Developmental Disorders at the University of Maryland, the MRC London Neurodegenerative Diseases Brain Bank, Kings College London, Denmark Hill, London, UK, and the Alzheimer’s Disease Sequencing Project (www.niagads.org/adsp for more details). Samples used in this research were obtained from the U.K. MND Collections funded by the MND Association and the Wellcome Trust. We thank Peter R. Schofield, William S. Brooks, Elizabeth M. Thompson, Peter Blumbergs, Cathy L. Short, Colin D. Field, Peter K. Panegyres, and Jane Hecker for their assistance in collecting samples from the Australian kindred.

## Author contributions

RC, RK, JEL, ACh, TC, BNS, and BJT designed the study, wrote the report, did the literature search, and drew the figures. OP, JT, DEM, The International ALS Genomics Consortium; The ITALSGEN Consortium; the FALS Consortium, Project MinE, IB, CD-S, JBK, RHB, ACa, GM, ACh, TC, CES, MG, BNS, and BJT obtained samples and clinical data from patients. JOJ, RC, RK, YA, FF, AER, HAP, SDT, NA, JRG, JD, MAN, CD-S, MN, CLD, SWS, MSS, SA, IG, FL, JEL, BNS, ACh, IJC, TC, and BJT performed experiments and the data analysis.

