## Supplementary Materials for "Mutations in the *SPTLC1* gene are a cause of juvenile amyotrophic lateral sclerosis that may be amenable to serine supplementation"

**Table of contents**


***Supplementary materials and methods***

- Clinical description of patient 1
- Clinical description of patient 2
- Clinical description of patient 3
- Immunohistochemistry
- Cellular mitochondrial assay
- Immunopurification of SPTLC1 protein complex
- Photometric serine palmitoyltransferase enzymatic assay
- Western blotting
- Data availability and software

***Supplementary figures***

- Figure S1. Biochemical assay of SPT activity and mitochondrial cellular phenotypes
- Figure S2. Distribution of *SPTLC1* mutations detected in adult-onset ALS patients.
- Figure S3. Chromatograms of *SPTLC1* variants identified in patients
- Figure S4. Immunohistochemistry of SPTLC1 in lumbar spinal cord
- Figure S5. Pedigrees of families carrying the p.Ala305Thr *SPTLC1* mutations

***Supplementary tables***

- Table S1. Clinical features of adult-onset ALS cases screened for *SPTLC1* mutations
- Table S2. Quantification of plasma sphingolipid concentrations using mass spectrometry
- Table S3. *SPTLC1* mutations identified in ALS cases
- Table S4. Primer sequences used for Sanger sequencing and mutagenesis of *SPTLC1*
- Table S5. The haplotype of chromosome 9q22.31 associated with the p.Ala305Thr variant in *SPTLC1*

***Other supplementary files***

- Movie S1. Tongue fasciculations and wasting in patient 2
- Movie S2. Gower's sign in patient 2
- Movie S3. Neurological manifestations in patient 3
- Movie S4. MRI brain and spinal cord in patient 2
- Consortia authors and affiliations

### Supplementary materials and methods

#### Detailed description of patient 1

Patient 1 is a right-handed, white woman who had presented at five years of age with spastic diplegia and motor delay. There was no family history of neurological disease, and her three siblings and her parents were neurologically healthy (see **Figure 2A**). By age 8, she had developed tongue fasciculations and proximal limb weakness. Her weight and height were consistently below the $10^{th}$ percentile despite treatment with growth hormones and oxandrolone. A gastrostomy tube was placed at age 10 due to anorexia.

Her muscle weakness progressed over the years. By age 19, she was unable to sit unsupported, stand or walk, and was entirely dependent for activities of daily living. Her hand weakness had progressed to the point that she could no longer write or use an iPad, and she had increasing difficulty using the joystick to control her motorized wheelchair. She complained of frequent and severe nocturnal leg spasms requiring treatment with valium, gabapentin, and trazodone.

She underwent tracheostomy at 17 years of age, and she has been on continuous ventilatory support since that time. The same year, she underwent spinal fusion surgery to correct scoliosis and lordosis that was causing restrictive lung disease, and right hip surgery for contractures. At age 18, she developed nighttime urinary urgency and intermittent bradycardia, likely secondary to increased vagal tone.

At age 16, the patient was referred to psychiatry for evaluation of depression, anxiety, and insomnia. At age 18, her mother noted that the patient was not as "witty and bright" as she had been. At age 20, the patient was having difficulty remembering things and was slow to speak in social settings.

Neurophysiology testing at ages 9, 10, and again at age 13 revealed diffuse chronic denervation, consistent with anterior horn cell dysfunction. Right sural nerve conduction amplitude and velocity were consistently reported to be normal. Cool detection threshold (CDT) and Q-Sweat testing, a commercial quantitative sweat measurement system, did not show evidence of small fiber sensory or autonomic dysfunction in distal peripheral nerves at the age of 20. Muscle biopsy of the left quadriceps at age 11 revealed chronic denervation and reinnervation with extensive atrophy involving entire fascicles and scattered large type 1 fibers.

Examination at the age of 20 revealed a short woman (122 cm, weight 28 kg, body mass index 18.8 $kg/m^2$, z-score = -1.12, $13^{th}$ percentile) sitting in a motorized wheelchair and ventilated via tracheostomy. The patient was dyspneic on talking, and she was moderately dysarthric. Her affect was jocular.

Montreal Cognitive Assessment yielded a score of 11 out of 25 due to poor orientation to date and location, poor serial sevens, impaired generation of words beginning with "F", poor abstraction, and poor delayed recall. Other visuospatial and executive functions were not assessed due to the patient's inability to write. Frontal Behavioral Inventory completed by the patient's mother gave a score of 7 [normal range: 0-27]. Frontotemporal Dementia Rating Scale, also completed by the patient's mother, identified a loss of interest, impulsivity, forgetfulness, and urinary and fecal incontinence (likely neurological in etiology, rather than arising from a behavioral issue).

Cranial nerve examination showed decreased upward eye gaze, bilateral fine nystagmus on lateral gaze, prominent perioral fasciculations, and a slow, wasted, fasciculating tongue. Palatal movements were also slow, and jaw jerk was absent. Shoulder shrug, neck flexion, and neck extension were weak (MRC = 3/5).

There was severe muscle wasting and weakness involving all four limbs. She had contractures of both feet with flexion of toes and foot inversion. Fasciculations were not observed. Ankle reflexes were brisk bilaterally, whereas knee jerks and upper limb reflexes were absent. Plantar responses were mute on Babinski testing (toe dorsiflexion and plantar flexion were 0/5). Pain sensation was decreased in the right arm and left leg. Temperature sensation, proprioception, and vibration sensation were normal.

**Detailed description of patient 2**

Patient 2 was a 15-year-old right-handed girl with mixed African American and white race/ethnicity who had developed difficulty with walking due to lower limb muscle weakness at the age of 8. By age 11, she had trouble climbing stairs. Tongue fasciculations were first observed at age 12. At the time of referral to her local medical center at the age of 15, she had generalized weakness and muscle wasting. She was unable to walk long distances without fatigue, and her weight was more than 7 standard deviations below average for her age. A prior endocrinology workup for her failure to gain weight was unremarkable. Her school performance had begun to decline at the age of 15.

Past medical history was otherwise unremarkable, and there was no family history of neurodegenerative disorders (**Figure 2B**).

Examination at the age of 15 revealed a thin girl with a body mass index below the 1$^{st}$ percentile based on her age and gender (z-score = -7). There was no evidence of orthostatic hypotension. Cranial nerve examination revealed tongue fasciculations and wasting (**Movie S1**).

Limb examination showed an exaggerated lumbar lordosis, generalized weakness involving all muscles, and prominent scapula winging bilaterally (**Figure 1A-B**). Her upper limb reflexes were diminished. In the lower limbs, knee jerks were reduced, and ankle jerks were asymmetrically brisk. Pinprick, temperature, vibration, and proprioception sensations were normal. The patient had an abnormal gait and a floridly positive Gower's sign (**Movie S2**).

MRI imaging of the brain and spine were unremarkable (**Movie S4**), though these imaging studies showed diffusely diminished pelvic and thigh musculature, and a notable lack of subcutaneous fat. EMG revealed active and chronic denervation with prominent complex repetitive discharges and giant muscle unit action potentials consistent with a long-standing motor neuropathic or neuronopathic disorder. There was no neurophysiological evidence for a sensory neuropathy. Plasma metanephrine and normetanephrine were normal. A neuropsychological evaluation revealed that her intellectual functioning was within the average range based on her age. However, she demonstrated significant challenges with sustained attention and stamina, aspects of executive functioning, and motor coordination. A swallow study was unremarkable.

**Detailed description of patient 3**

Patient 3 was an eleven-year-old African American right-handed girl with a history of failure to gain weight and toe-walking since the age of four. She was first evaluated at the age of 10 because of difficulty walking due to her left foot "turning in." In retrospect, her gait had been gradually deteriorating over several years, resulting in occasional falls and difficulty walking upstairs. She was no longer able to participate in her school dance class due to weakness and exercise-induced cramps. Subsequently, her symptoms had spread to her hands. She was unable to undo soda bottle tops, and dressing has been taking longer than usual. The patient had complained of tingling in her feet over the last three years, but the numbness was only noticed on her first neurological evaluation. She had a painless ulcer on her right foot that developed from her ankle-foot orthosis. She has occasional choking episodes, mostly on liquids. She has noticed the pooling of secretions. Her speech has been normal, and there have been no obvious cognitive or behavioral problems. She has worn glasses since age eight. Ophthalmological examination at age eleven revealed bilateral cataracts that did not require surgical intervention. The patient has a history of profuse sweating, but no urinary symptoms or dizziness on standing.

Her past medical history included adenoidectomy, bilateral Achilles tenotomy at the age of 10, and long-standing vitamin D deficiency requiring oral supplementation. She was born one month premature, but she achieved her early developmental milestones appropriately. There was no family history of ALS, and her nineteen-year-old brother was healthy (**Figure 2C**). Her maternal grandmother, paternal grandfather, and maternal aunt were diagnosed with late-onset dementia.

Examination at the age of eleven revealed a thin girl with a body mass index of 10.9 kg/m2, representing less than the 1$^{st}$ percentile based on her age and gender (z score = -6.5). Blood pressure was 111/62 mmHg with a heart rate of 107 beats per minute while lying, and 128/62 mmHg with a heart rate of 139 beats per minute on standing. She was alert and articulate, and her conversational speech was normal, though she had difficulty forming K and L sounds.

On cranial nerve examination, her near visual acuity was 20/50 bilaterally without glasses, and fundoscopy confirmed the presence of lens opacities. The tongue was wasted, weak, and slow to move with prominent fasciculations (**Figure 1C-D**). Jaw jerk was present, and shoulder shrug was weak bilaterally. The rest of the cranial nerves were intact.

Her posture was normal, and there was no evidence of scoliosis. The examination of her limbs showed generalized atrophy and hypotonia. Fasciculations were present in the upper and lower limbs. There was a left greater than right pronator drift with bilateral postural tremor. The patient had bilateral hammertoes, high arches, and a healing ulcer over the right first metatarsal joint. Strength was decreased in a pyramidal pattern that was asymmetric and affecting distal muscles more than proximal. Power in the upper limbs was as follows: bilateral shoulder abductors = 4/5, shoulder adductors = 5/5, right elbow flexor = 5/5, left elbow flexor = 4/5, bilateral elbow extensors = 4/5, wrist dorsiflexor = 4/5, wrist palmar flexion = 5/5, finger flexors = 4/5, right finger extensors = 4/5, left finger extensors = 3/5, bilateral finger adductors = 2/5, finger abductors = 3/5, and thumb abductors = 4/5. Power in the lower limbs was as follows: bilateral hip flexors = 4/5, hip abductors = 4/5, hip adductors = 5/5, knee flexors = 4/5, knee extensors = 5/5, ankle dorsiflexion = 3/5, ankle plantar flexion = 5/5, ankle eversion = 3/5, ankle inversion = 5/5, hallux flexors = 2/5, and hallux extensors = 2/5. Reflexes in the upper limbs were 4+ with spread with a positive Hoffman's sign bilaterally. In the lower limbs, knee jerks were 3+ bilaterally, and ankle jerks were absent. The left Babinski was down going, whereas the right Babinski elicited fanning of her toes. The patient was ataxic on heel-to-shin and finger-to-nose testing with bilateral dysdiadochokinesis, though this appeared to be secondary to muscle weakness. The sensory exam revealed decreased pinprick and temperature sensation in the upper limbs to the level of the wrist, right worse than left, and in the lower limbs to the mid-calf level. Proprioception and vibration sensations were intact. The patient walked abnormally due to generalized weakness and bilateral foot drop (**Movie S2**). She had difficulty with tandem gait testing, was able to stand on her toes, but she was unable to stand on her heels. Romberg's sign was negative. She had a positive Gower's sign.

Neurophysiological examination at the age of 11 was consistent with sensorimotor axonal neuropathy. Electromyography of the left vastus lateralis and tibialis anterior muscles showed large amplitude, polyphasic potentials with reduced amplitude pattern. MRI of the brain and spinal cord was unremarkable. Pulmonary function testing demonstrated a mild restrictive ventilatory defect (forced expiratory volume in one second-to-forced vital capacity ratio = 0.9). Creatine phosphokinase, vitamin B12, and folate serum transferrin, iron, copper, metabolic screens were within normal limits. Serine level was within the normal range at 79.3 umol/l.

**Recruitment**
Patients 1, 2, and 3 were followed by their local neurologists and geneticists. All patients were diagnosed with juvenile ALS by their local physicians. Patients 1 (and her family) was referred to Dr. Traynor at the NIH Clinical Center for evaluation by Dr. Crawford, her pediatric neurologist at Hopkins University. Dr. Traynor has a joint appointment at Hopkins University. Dr. Traynor was contacted by the physicians of patients 2 and 3 through the GeneMatcher program at GeneDx (*35*). Patient 2 declined referral to the NIH Clinical Center. Patient 3 (and her family) was referred to Dr. Traynor at the NIH Clinical Center for evaluation by Dr. Li, her pediatrician at Virginia Commonwealth University. The NIH and Virginia Commonwealth University are regional partners in research based on a formal master affiliation agreement between the two institutions.

**Immunohistochemistry**
The control spinal cord was obtained from the postmortem of a 64-year-old white man, and the postmortem interval was 28 hours. The sporadic ALS spinal cord was obtained from the postmortem of a 66-year old white male, and the postmortem interval was 4.5 hours. The spinal cord carrying the *SPTLC1* p.Arg445Gln mutation was obtained from the postmortem of a 78-year-old white man, who had been diagnosed with sporadic ALS. Postmortem interval was 3.83 hours, and survival from symptom onset in his upper limbs was 68 months. The autopsy of the *SPTLC1* carrier was performed according to the research protocol of the Department of Veterans Affairs Biorepository Brain Bank (*35*). The other autopsies were performed at the JHMI Brain Resource Center according to their internal protocol.

Formalin-fixed, paraffin-embedded blocks were sectioned to 5 um thickness and stained with primary anti-SPTLC1 antibody (Sigma-Aldrich) at 1:50 dilution overnight at 4°C and counter-stained with hematoxylin.

#### Cellular mitochondrial assay

Mutations were introduced into a plasmid containing the human *SPTLC1* open reading frame (Origene) using the QuikChange II XL kit (Agilent), followed by subcloning into pLenti-C-Myc-DDK-P2A-Puro lentiviral plasmid (Origene). Lentiviruses were produced with third-generation packaging plasmids (pMDLg/pRRE and pRSV-Rev, Addgene) and envelope plasmid (pMD2.G, Addgene) (*36*). HEK293FT were transfected with wild-type or mutant lentivirus transfer plasmid, and transduced cells stably expressing SPTLC1 were selected by extended growth in 0.5 ug/ml puromycin (Thermo Fisher). For the serine rescue experiment, a final concentration of 100mM L-serine was added to the media for 48 hours. Mitochondria in live cells were incubated with 100 nM MitoTracker Red CMXRos (Thermo Fisher) at 37°C for one hour, then fixed with 4% paraformaldehyde, followed by washing with PBS before imaging using a high-content imager (CellInsight, Thermo Fisher). A minimum of six wells were quantified for each condition, and all assays were performed at least twice. Unpaired t-test with Welch's correction or ANOVA were used to calculate statistical significance.

#### Immunopurification of SPTLC1 protein complex

SPTLC1-FLAG proteins were immuno-purified from HEK293FT cells that were stably expressing SPTLC1. Cells were lysed in lysis buffer (50 mM HEPES, pH8.0, 1 mM EDTA, 0.1% (w/v) sodium monolaurate, phosphatase (Thermo Fisher), and protease inhibitor (Roche AG)). Membrane proteins were further solubilized by sonication for fifteen seconds at 50% power and 50% pulsation for a total of 45 seconds. The lysate was clarified by centrifugation at 21,000g for five minutes at 4°C to remove large cellular debris followed by immunoprecipitation with EZview Red Anti-Flag M2 Affinity Gel (Sigma-Aldrich) as previously described (*37*). Purified SPTLC1-FLAG was quantified against a bovine serum albumin protein standard curve (BSA, Thermo Fisher) ran on an SDS-PAGE gel and transferred to a nitrocellulose membrane (Bio-rad). The membrane was stained with Revert Total Protein Stain (LI-COR) and imaged on an Odyssey CLx imaging system. Interaction of SPTLC2 with SPTLC1-FLAG was confirmed by Western blot showing that the purified serine palmitoyltransferase complex was likely functional.

#### Photometric serine palmitoyltransferase enzymatic assay

Condensation of serine and palmitoyl-CoA by serine palmitoyltransferase enzyme produces 3-ketodihydrosphingosine and releases both carbon dioxide and free coenzyme A (CoA). This photometric assay measures the amount of free CoA released that is reactive to 5,5'-dithiobis-2-nitrobenzoic acid (DTNB, Thermo Fisher) which is reflective of the enzymatic activity of serine palmitoyltransferase.

This assay was adapted from and shown to be comparable to radioactive assays measuring radiolabeled 3-ketodihydrosphingosine (*38*). Triplicate wells were assayed per 100ng of purified wild-type or mutant SPTLC1 proteins, and per amino acid tested. The product of the reaction was measured at 412 nm at the zero-time point and every two minutes for up to one hour. In place of purified SPTLC1-FLAG protein, varying concentrations of CoA, were included to build the calibration curve for the estimation of CoA released from the serine palmitoyltransferase enzymatic reaction. Estimated amounts of CoA produced per nanogram of SPTLC1-FLAG protein were plotted over time for each amino acid.

#### Western blotting

Western blot was performed as previously described (*37*). We used the following primary antibodies at the indicated dilutions: mouse anti-Flag (1:5,000, Sigma-Aldrich), rabbit anti-SPTLC1 (1:1,000, Sigma-Aldrich), and rabbit anti-SPTLC2 (1:1000, LifeSpan BioSciences Inc.).

#### Data and materials availability

Alzheimer's Disease Sequencing Project (ADSP): https://www.niagads.org/adsp/content/home. Database of Genotypes and Phenotypes (dbGaP): https://www.ncbi.nlm.nih.gov/gap. Genome Aggregation Database (gnomAD):

https://gnomad.broadinstitute.org. Haplotype Reference Consortium (HRC): www.haplotype-reference-consortium.org. Kaviar Genomic Variant Database: http://db.systemsbiology.net/kaviar/. ANNOVAR: annovar.openbioinformatics.org. Clustal Omega: www.ebi.ac.uk/Tools/msa/clustalo/. GATK: http://www.broadinstitute.org/gsa/wiki/index.php/Home_Page. MassLynx: http://www.waters.com/waters/en_US/MassLynx-Mass-Spectrometry-Software-/nav.htm?cid=513164&locale=en_US. Picard: http://broadinstitute.github.io/picard. PLINK: https://www.cog-genomics.org/plink/1.9/. Sequencher: http://genecodes.com. TRAPD: https://github.com/mhguo1/TRAPD.

### Supplementary Figures

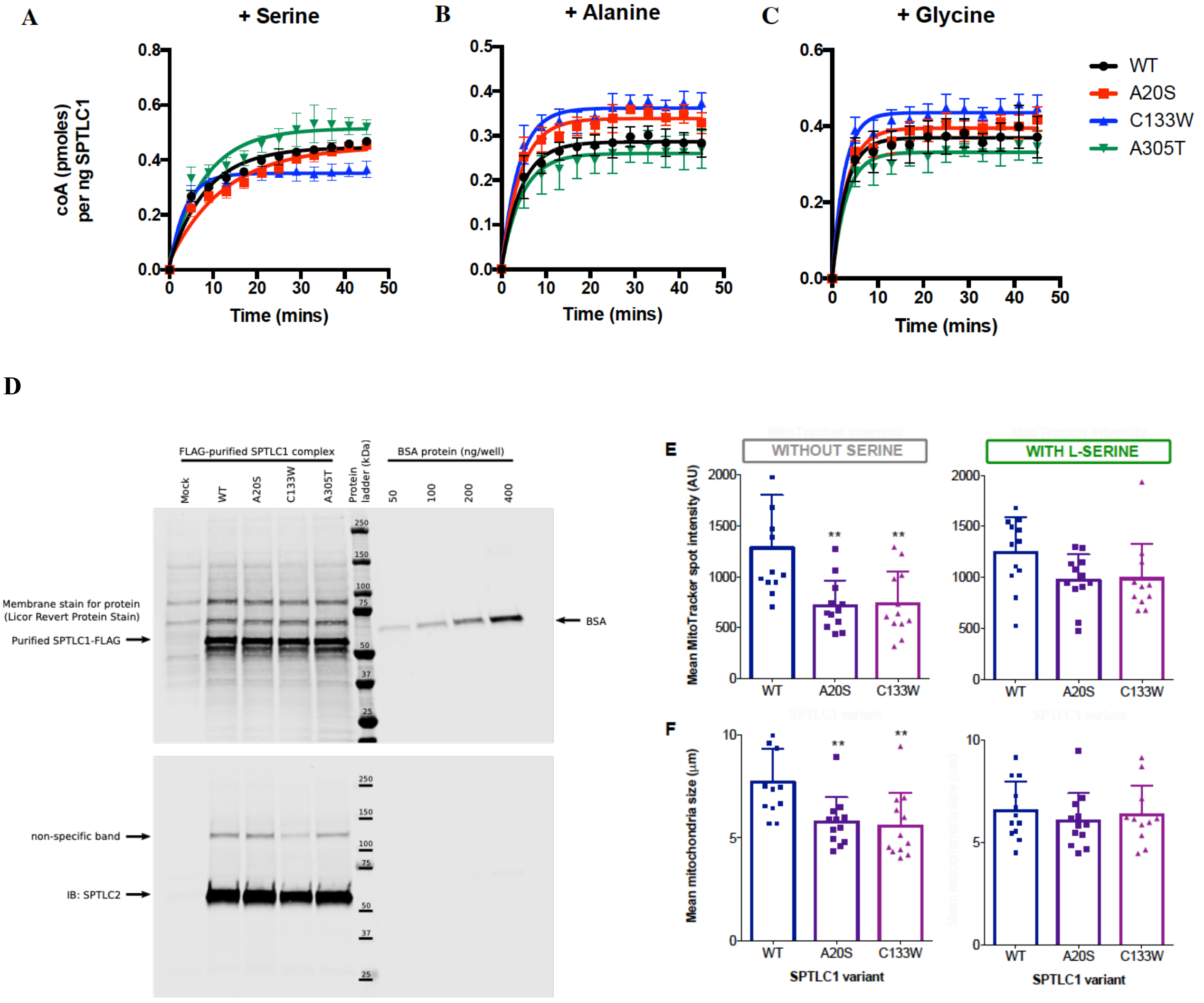


**Figure S1. Biochemical assay of SPT activity and mitochondrial cellular phenotypes.** The activity of purified SPTLC1 enzyme complex was determined using a photometric assay by measuring the release of free coA from the condensation reaction between palmitoyl-CoA and **(A)** L-serine, **(B)** L-alanine and **(C)** L-glycine. All wild-type and mutant SPTLC1 complexes (p.Ala20Ser, p.Cys133Trp, and p.Ala305Thr) are active enzymes capable of utilizing all three amino acids as substrates. The notable difference between the mutant SPTLC1 complex (p.Ala20Ser and p.Cys133Trp) is the altered preference for L-alanine or glycine over L-serine, compared to the wild-type SPTLC1 complex. **(D)** Western blot of purified SPTLC1-FLAG proteins showing that the purified complex contains SPTLC2**.** Mitochondria in HEK293 cells expressing wild type (WT), p.Ala20Ser, and p.Cys133Trp were assessed using MitoTracker on a high-content imager. **(E)** Mitochondrial intensity and **(F)** mitochondria size were smaller in cells expressing mutant protein under standard culture conditions. Supplementation of 100 mM L-serine in culture media for 48 hours rescued the mitochondrial abnormalities in the p.Ala20Ser and p.Cys133Trp lines **(E, F)**.

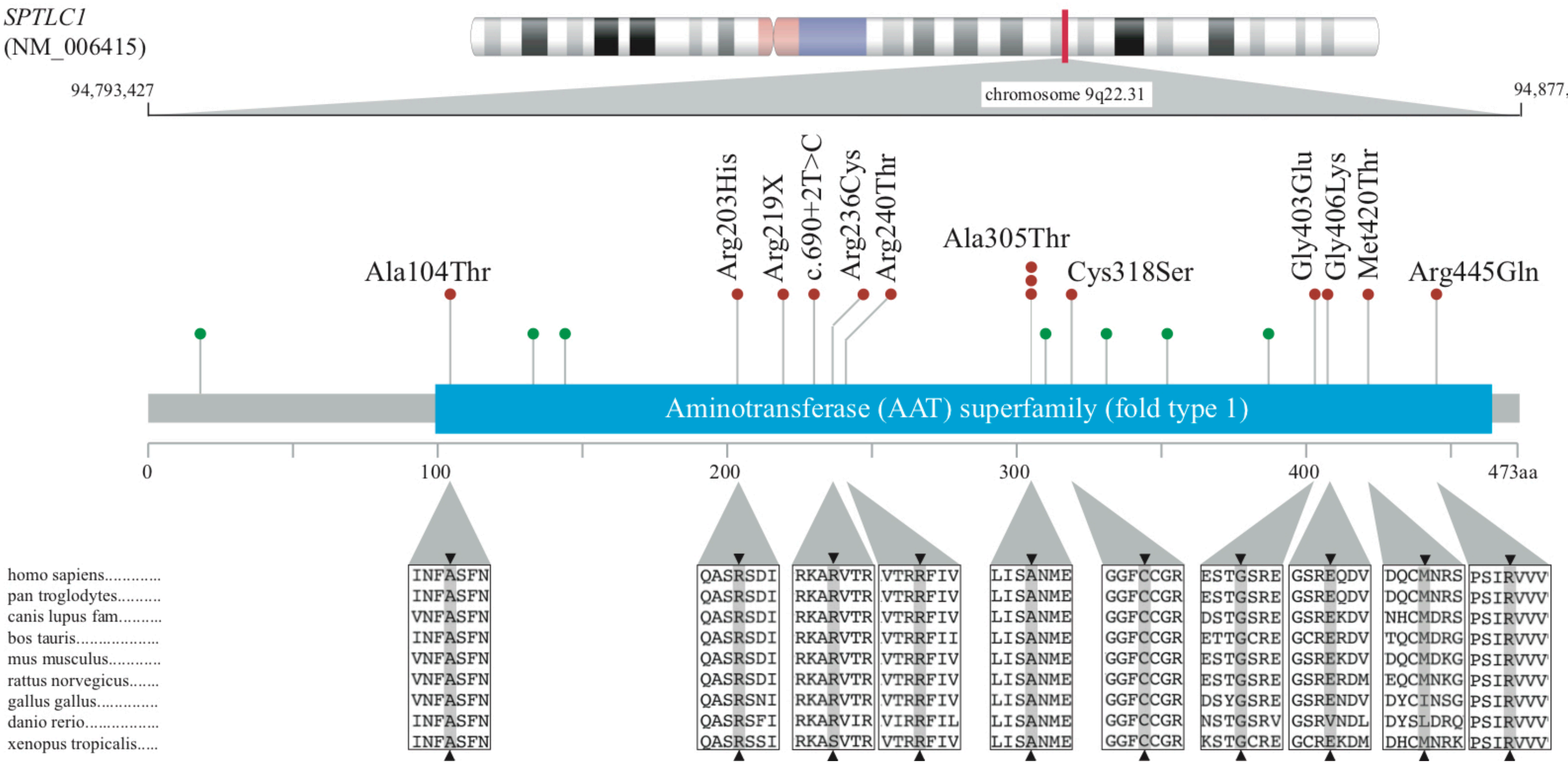


**Figure S2. Distribution of SPTLC1 mutations detected in adult-onset ALS patients.**
The top panel is a graphical representation showing the domains of SPTLC1. Mutations detected in ALS cases are indicated in red, and mutations previously described to cause HSAN1 are shown in green. The p.Ala305Thr variant was found in three ALS patients, and the other variants were found in single cases of adult-onset ALS. The middle panel shows the conservation of amino acid residue across species generated using the Clustal Omega online tool. The bottom panel portrays population frequency data and functional predictions of the mutations. A red filled square means that a variant met criteria.

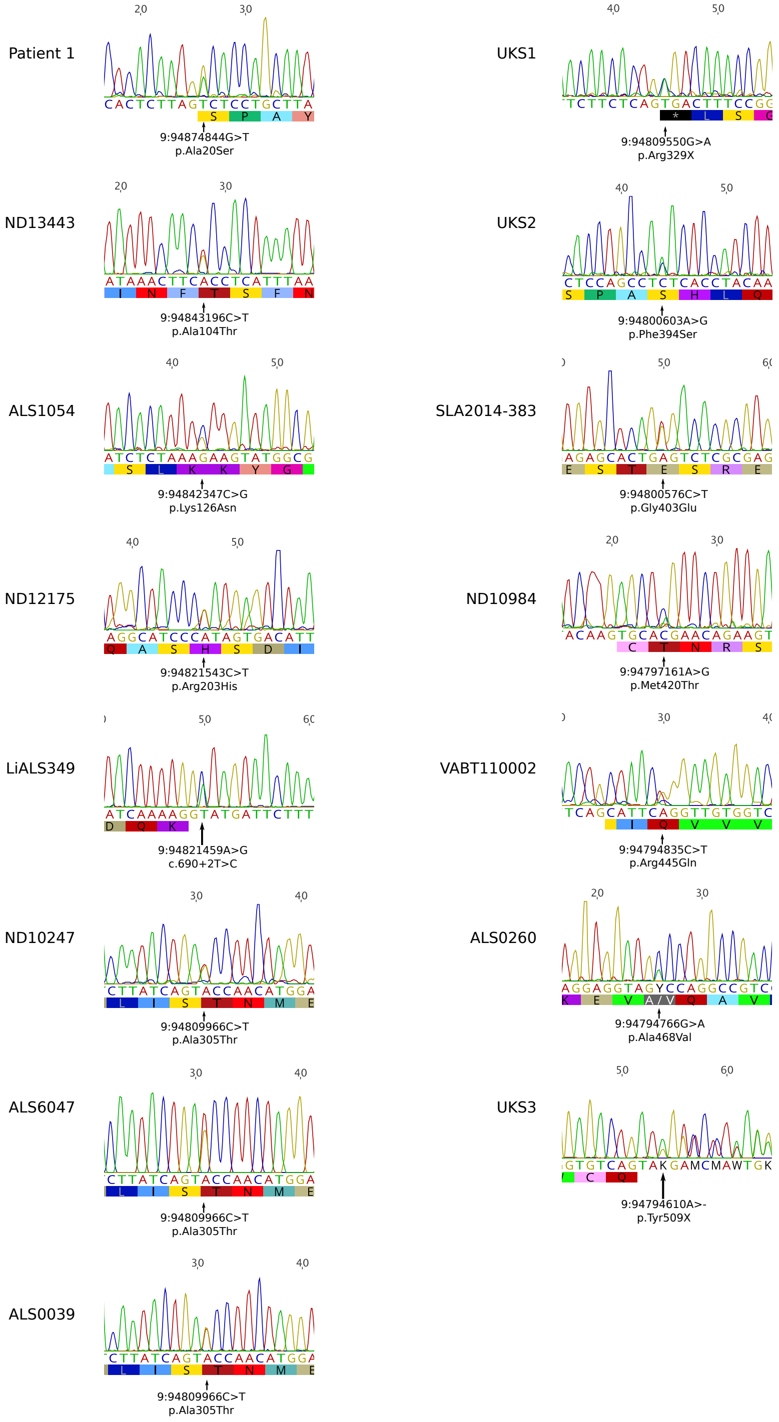


**Figure S3. Chromatograms of *SPTLC1* variants identified in patients diagnosed with juvenile ALS and adult-onset ALS.**

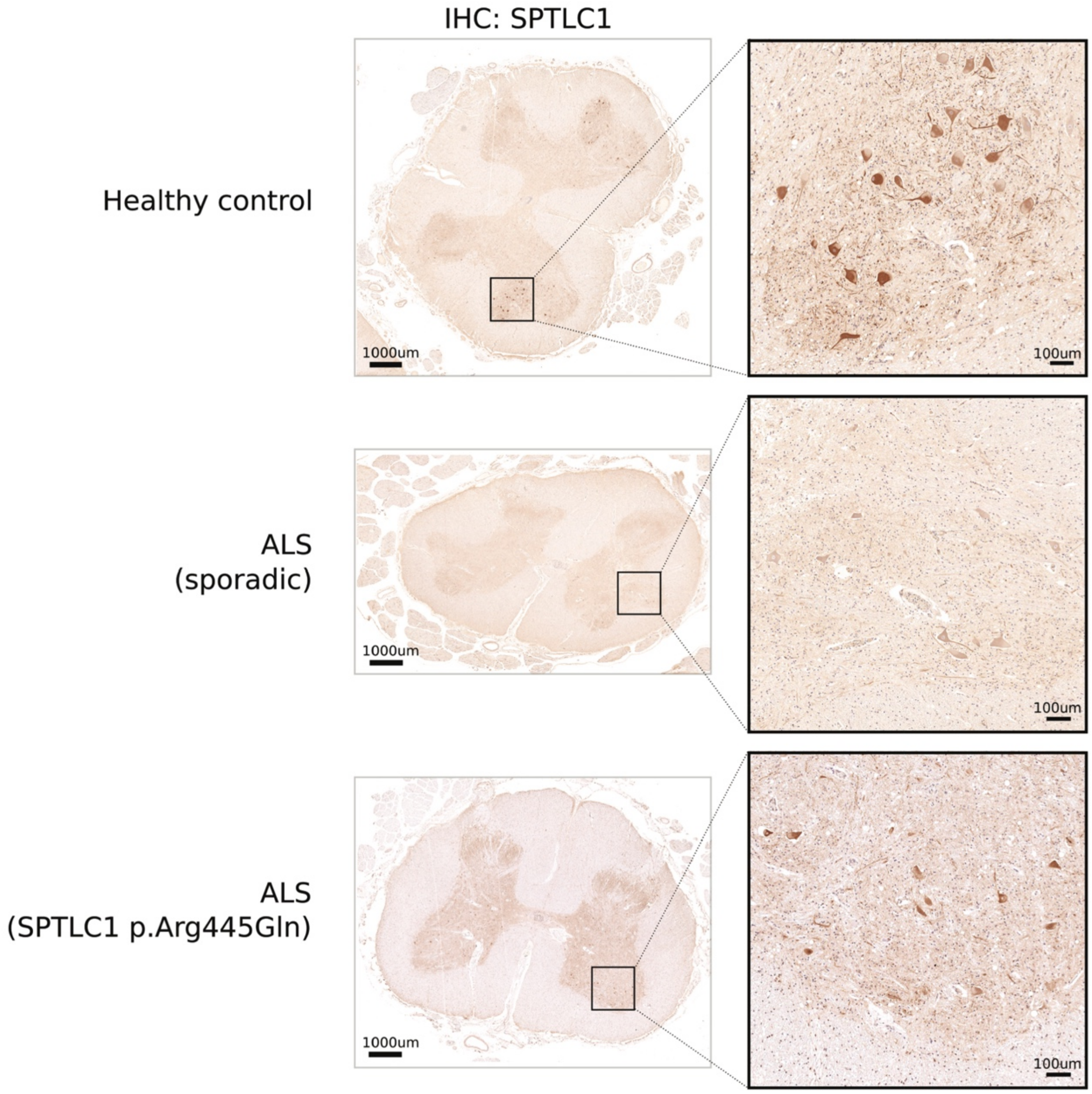


**Figure S4. Immunohistochemistry of SPTLC1 in the lumbar spinal cord.**
The control spinal cord (top panel) exhibits SPLTC1 immunoreactivity in the lamina IX motor neurons of the anterior horns. The spinal cord from a subject with sporadic ALS (middle panel) shows weaker immunoreactivity in the same region, consistent with a decrease in the number and size of surviving motor neurons. The spinal cord from an ALS case carrying the p.Arg445Gln mutation in SPTLC1 (bottom panel) shows a similar pattern of diminished staining as the sporadic ALS sample. Scale bars in magnified inset represent 100 μm.

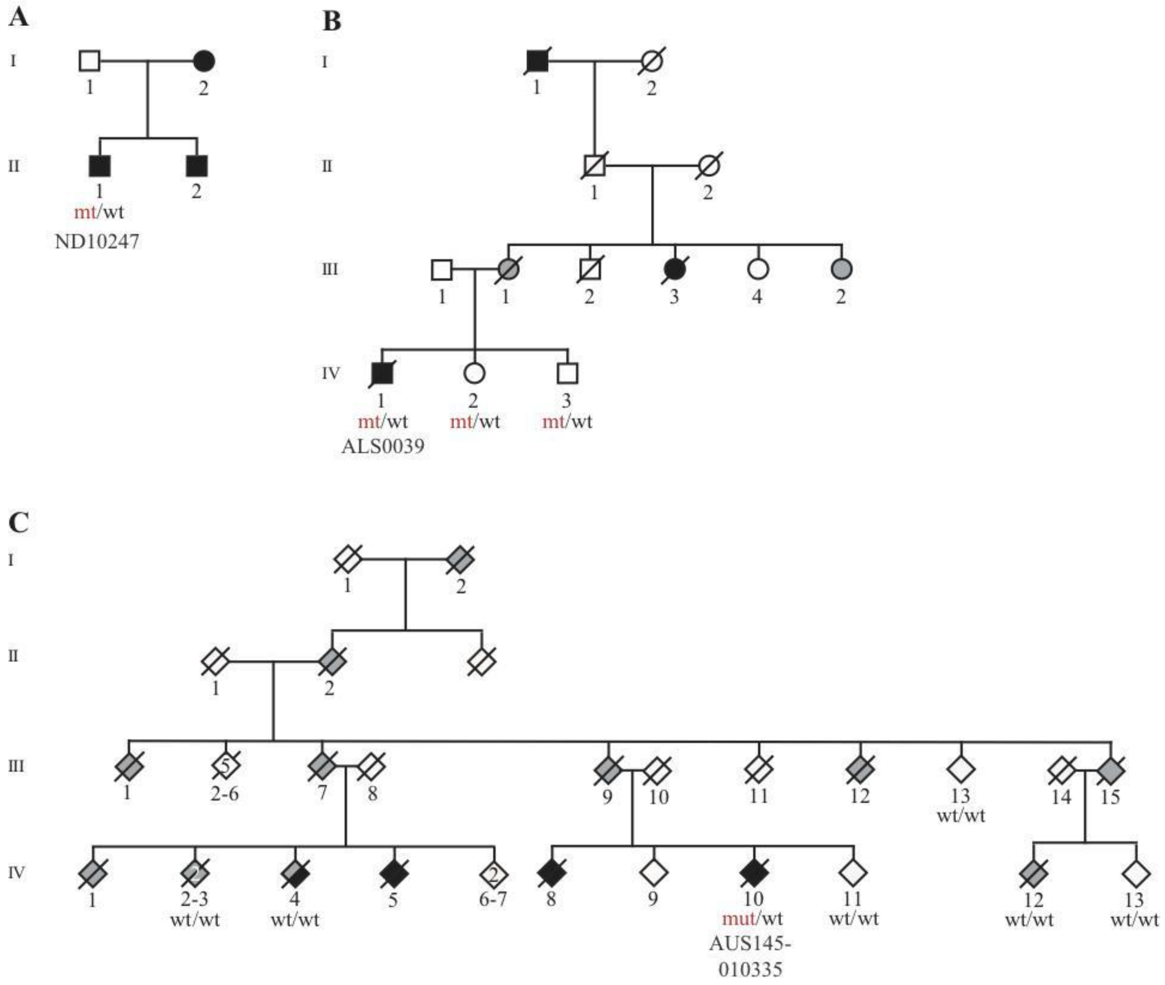


**Figure S5. Pedigrees of families carrying the p.Ala305Thr *SPTLC1* mutations**
Representations of the British **(A)**, the American **(B),** and the Australian **(C)** kindreds carrying the p.Ala305Thr mutation. The mutant allele is indicated by *mt*, whereas wild-type alleles are indicated by *wt*. Shapes filled with black represent individuals diagnosed with ALS, and forms filled with grey represent individuals diagnosed with FTD.

### Supplementary Tables

**Table S1. Clinical features of adult-onset ALS cases screened for *SPTLC1* mutations**

| | **ALS cases (n = 5607)** | **Control subjects (n=5,710)** |
|---|---|---|
| Mean age at onset/sampling (SD) | 57.4 (14.9) | 86.1 (6.9) |
| Female (%) | 2,017 (42.7%) | 3,370 (59.0%) |
| Site of onset: | | |
| Bulbar (%) | 979 (26.7%) | NA |
| Spinal (%) | 2,693 (73.3%) | NA |
| Familial disease (%) | 2,322 (46.1%) | NA |

ALS cases include samples from the USA (n = 1,922), Italy (n = 1,295), Finland (n = 306), Germany (n = 243), United Kingdom (n = 851) and other countries (n = 128); Control samples include samples from the USA (n = 4,647), Italy (n = 763) and Finland (n = 300); Control participants consisted of samples that had undergone whole-genome sequencing (n = 300 Finnish and n = 677 Italian) and samples that had undergone exome sequencing as part of the Alzheimer's Disease Sequencing Project (n = 4,647 Americans). Clinical data was missing for age at onset (n = 1,707), sex (n = 882), site of onset (n = 1,935), family status (n = 567) and country (n = 704).

**Table S2. Quantification of plasma sphingolipid concentrations using mass spectrometry**

**Compound 1: 1-deoxy methyl sphingosine**

| Sample | Std. Conc | RT | Area | IS Area | Response | Conc. (ng/ul) | %Dev | S/N |
|---|---|---|---|---|---|---|---|---|
| Solvent blank | | 3.67 | 21139.021 | 234224.328 | 0.09 | 4.8 | | 26.403 |
| 1 ng | 1 | 3.69 | 6929.532 | 128369.969 | 0.054 | 1.1 | 14.5 | 6.301 |
| 10 ng | 10 | 3.69 | 14549.364 | 115930.992 | 0.126 | 8.3 | -17 | 10.221 |
| 25 ng | 25 | 3.68 | 34235.965 | 125188.125 | 0.273 | 23.1 | -7.6 | 20.044 |
| 100 ng | 100 | 3.67 | 91720.023 | 88701.75 | 1.034 | 99.1 | -0.9 | 28.518 |
| 500 ng | 500 | 3.67 | 225606.969 | 42056.617 | 5.364 | 532.2 | 6.4 | 36.955 |
| 1000 ng | 1000 | 3.65 | 291422.594 | 27122.564 | 10.745 | 1070.2 | 7 | 47.183 |
| 2000 ng | 2000 | 3.65 | 366156.625 | 17825.775 | 20.541 | 2049.8 | 2.5 | 49.688 |
| 3000 ng | 3000 | 3.65 | 425962.125 | 14911.81 | 28.565 | 2852.2 | -4.9 | 46.961 |
| 4000 ng | 4000 | 3.65 | 466330.625 | 12867.295 | 36.242 | 3619.8 | -9.5 | 45.581 |
| blank | | 4.1 | 7767.99 | 293.278 | 26.487 | 2644.3 | | 1.222 |
| blank | | | | 2144.102 | | | | |
| QC-1 | 750 | 3.66 | 274286.594 | 49900.813 | 5.497 | 545.4 | -27.3 | 159.211 |
| QC-2 | 1500 | 3.65 | 369979.438 | 32661.219 | 11.328 | 1128.5 | -24.8 | 122.384 |
| QC-3 | 2500 | 3.65 | 415159.719 | 20376.768 | 20.374 | 2033.1 | -18.7 | 87.671 |
| blank | | 3.68 | 4558.362 | | | | | 4.221 |
| blank | | | | 454.999 | | | | |
| blank | | | | 658.992 | | | | |
| QC-2 | 1500 | 3.66 | 346855.781 | 30974.672 | 11.198 | 1115.5 | -25.6 | 120.801 |
| blank | | | | 707.815 | | | | |
| blank | | | | 334.798 | | | | |
| LNG_P1 (unrelated pop. control) | | 3.71 | 13976.459 | 146764.578 | 0.095 | 5.3 | | 5.374 |
| LNG_P2 (unrelated pop. control) | | 3.71 | 9069.832 | 140827.719 | 0.064 | 2.2 | | 7.238 |
| 2013-303 (Mother - unaffected) | | 3.7 | 13819.86 | 146548.563 | 0.094 | 5.2 | | 8.62 |
| 2013-302 (Proband) | | 3.7 | 14781.934 | 131780.281 | 0.112 | 7 | | 10.176 |
| 2013-307 (Brother - unaffected) | | 3.7 | 8705.323 | 153177.344 | 0.057 | 1.4 | | 8.125 |
| blank | | | | 707.815 | | | | |
| blank | | | | 334.798 | | | | |

**Compound 2: 1-deoxy methyl sphinganine**

| Sample | Std. Conc | RT | Area | IS Area | Response | Conc. | %Dev | S/N |
|---|---|---|---|---|---|---|---|---|
| blank | | 3.68 | 1847.313 | | | | | 1.404 |
| blank | | 3.69 | 411.352 | | | | | 0.641 |
| Solvent blank | | 3.69 | 218384.313 | 234224.328 | 0.932 | 20.6 | | 484.13 |
| 1 ng | 1 | 3.69 | 81351.25 | 128369.969 | 0.634 | 1.2 | 18.7 | 194.064 |
| 10 ng | 10 | 3.69 | 86499.336 | 115930.992 | 0.746 | 8.5 | -15 | 174.64 |
| 25 ng | 25 | 3.69 | 118198.891 | 125188.125 | 0.944 | 21.4 | -14.4 | 259.111 |
| 100 ng | 100 | 3.69 | 202974.266 | 88701.75 | 2.288 | 110.1 | 10.1 | 469.235 |
| 500 ng | 500 | 3.69 | 340958.656 | 42056.617 | 8.107 | 518.3 | 3.7 | 572.438 |
| 1000 ng | 1000 | 3.69 | 374393.563 | 27122.564 | 13.804 | 967.3 | -3.3 | 272.345 |
| 2000 ng | 2000 | 3.7 | 371646.5 | 17825.775 | 20.849 | 1627.7 | -18.6 | 127.131 |
| 3000 ng | 3000 | 3.71 | 404361.906 | 14911.81 | 27.117 | 2407.2 | -19.8 | 76.915 |
| 4000 ng | 4000 | 3.71 | 425882.313 | 12867.295 | 33.098 | 4098.3 | 2.5 | 63.395 |
| blank | | 3.69 | 12181.052 | 293.278 | 41.534 | | | 29.707 |
| blank | | 3.69 | 3663.977 | 2144.102 | 1.709 | 71.6 | | 12.778 |
| QC-1 | 750 | 3.7 | 393027.719 | 49900.813 | 7.876 | 501.3 | -33.2 | 317.144 |
| QC-2 | 1500 | 3.7 | 409117 | 32661.219 | 12.526 | 861.4 | -42.6 | 138.456 |
| QC-3 | 2500 | 3.7 | 419644.5 | 20376.768 | 20.594 | 1601 | -36 | 87.006 |
| blank | | 3.69 | 8047.265 | | | | | 24.414 |
| blank | | 3.69 | 3371.57 | 454.999 | 7.41 | 467.1 | | 9.751 |
| blank | | 3.69 | 620.522 | 658.992 | 0.942 | 21.2 | | 0.786 |
| QC-2 | 1500 | 3.7 | 402007.375 | 30974.672 | 12.979 | 898.5 | -40.1 | 129.135 |
| blank | | 3.69 | 505.912 | 707.815 | 0.715 | 6.5 | | 0.92 |
| blank | | 3.62 | 1160.128 | 334.798 | 3.465 | 189.3 | | 0.732 |
| LNG_P1 (unrelated pop. control) | | 3.71 | 109217.953 | 146764.578 | 0.744 | 8.4 | | 133.482 |
| LNG_P2 (unrelated pop. control) | | 3.7 | 114331.102 | 140827.719 | 0.812 | 12.8 | | 240.863 |
| 2013-303 (Mother - unaffected) | | 3.7 | 120275.234 | 146548.563 | 0.821 | 13.4 | | 336.212 |
| 2013-302 (Proband) | | 3.7 | 122818.75 | 131780.281 | 0.932 | 20.6 | | 367.057 |
| 2013-307 (Brother - unaffected) | | 3.7 | 118922.445 | 153177.344 | 0.776 | 10.5 | | 212.996 |
| blank | | 3.69 | 505.912 | 707.815 | 0.715 | 6.5 | | 0.92 |
| blank | | 3.62 | 1160.128 | 334.798 | 3.465 | 189.3 | | 0.732 |

**Compound 3: 1- deoxy sphingosine**

| Sample | Std. Conc | RT | Area | IS Area | Response | Conc. | %Dev | S/N |
|---|---|---|---|---|---|---|---|---|
| blank | | 3.64 | 6038.714 | 8036.412 | 0.751 | 121.4 | | 5.833 |
| blank | | 3.65 | 1788.422 | 2846.76 | 0.628 | 101.3 | | 2.018 |
| Solvent blank | | 3.68 | 58985.793 | 568738.125 | 0.104 | 15.5 | | 127.808 |
| 1 ng | 1 | 3.66 | 3467.167 | 240637.609 | 0.014 | 0.9 | -9.9 | 6.36 |
| 10 ng | 10 | 3.68 | 12637.877 | 231276.203 | 0.055 | 7.5 | -25.2 | 22.697 |
| 25 ng | 25 | 3.69 | 61421.738 | 240532.516 | 0.255 | 40.3 | 61.2 | 102.578 |
| 100 ng | 100 | 3.69 | 195372.781 | 195340.188 | 1 | 162.1 | 62.1 | 431.461 |
| 500 ng | 500 | 3.69 | 480828.25 | 127164.383 | 3.781 | 616.8 | 23.4 | 563.202 |
| 1000 ng | 1000 | 3.68 | 640941.375 | 86727.977 | 7.39 | 1207 | 20.7 | 461.563 |
| 2000 ng | 2000 | 3.68 | 841757.5 | 68812.102 | 12.233 | 1998.8 | -0.1 | 285.441 |
| 3000 ng | 3000 | 3.68 | 1047744.5 | 59297.875 | 17.669 | 2887.7 | -3.7 | 224.546 |
| 4000 ng | 4000 | 3.68 | 1098994 | 47366.727 | 23.202 | 3792.4 | -5.2 | 219.685 |
| blank | | 3.68 | 10682.125 | 3169.399 | 3.37 | 549.7 | | 20.338 |
| blank | | 3.68 | 4945.147 | 2604.694 | 1.899 | 309 | | 9.794 |
| QC-1 | 750 | 3.68 | 570821.188 | 273212.469 | 2.089 | 340.2 | -54.6 | 497.565 |
| QC-2 | 1500 | 3.68 | 830184.875 | 214402.469 | 3.872 | 631.7 | -57.9 | 304.198 |
| QC-3 | 2500 | 3.68 | 980890.688 | 138335 | 7.091 | 1158 | -53.7 | 258.314 |
| blank | | 3.69 | 8024.52 | 3600.424 | 2.229 | 363 | | 12.675 |
| blank | | 3.69 | 3661.554 | 3254.976 | 1.125 | 182.5 | | 6.309 |
| blank | | 3.67 | 1918.787 | 2818.832 | 0.681 | 109.8 | | 2.802 |
| QC-2 | 1500 | 3.69 | 822858.813 | 205209 | 4.01 | 654.2 | -56.4 | 304.143 |
| blank | | 3.67 | 1810.653 | 3074.438 | 0.589 | 94.8 | | 2.47 |
| blank | | 3.66 | 1851.128 | 2633.729 | 0.703 | 113.5 | | 2.302 |
| LNG_P1<br>(unrelated pop. control) | | 3.66 | 3155.857 | 335556.094 | 0.009 | 0.1 | | 5.186 |
| LNG_P2<br>(unrelated pop. control) | | 3.67 | 2450.698 | 334425.938 | 0.007 | 0 | | 5.866 |
| 2013-303<br>(Mother - unaffected) | | 3.66 | 2466.829 | 340491.313 | 0.007 | 0 | | 6.294 |
| 2013-302<br>(Proband) | | 3.67 | 2640.166 | 354765.781 | 0.007 | 0 | | 4.245 |
| 2013-307<br>(Brother - unaffected) | | 3.67 | 2552.419 | 343583.031 | 0.007 | 0 | | 4.16 |
| blank | | 3.67 | 1810.653 | 3074.438 | 0.589 | 94.8 | | 2.47 |
| blank | | 3.66 | 1851.128 | 2633.729 | 0.703 | 113.5 | | 2.302 |

**Compound 4: 1-deoxy sphinganine**

| Sample | Std. Conc | RT | Area | IS Area | Response | Conc. | %Dev | S/N |
|---|---|---|---|---|---|---|---|---|
| blank | | 3.56 | 4651.65 | 8036.412 | 0.579 | 345.2 | | 10.127 |
| blank | | 3.57 | 1188.88 | 2846.76 | 0.418 | 242.7 | | 3.719 |
| Solvent blank | | 3.7 | 11875.195 | 568738.125 | 0.021 | 2.6 | | 42.878 |
| 1 ng | 1 | 3.62 | 4302.635 | 240637.609 | 0.018 | 0.9 | -12.5 | 21.009 |
| 10 ng | 10 | 3.62 | 4920.925 | 231276.203 | 0.021 | 2.9 | -71.4 | 10.677 |
| 25 ng | 25 | 3.7 | 14562.527 | 240532.516 | 0.061 | 25.9 | 3.6 | 36.258 |
| 100 ng | 100 | 3.7 | 39127.426 | 195340.188 | 0.2 | 109.2 | 9.2 | 136.464 |
| 500 ng | 500 | 3.7 | 106171.227 | 127164.383 | 0.835 | 515 | 3 | 412.898 |
| 1000 ng | 1000 | 3.7 | 125247.063 | 86727.977 | 1.444 | 962 | -3.8 | 489.258 |
| 2000 ng | 2000 | 3.71 | 136210.656 | 68812.102 | 1.979 | 1428.3 | -28.6 | 216.563 |
| 3000 ng | 3000 | 3.72 | 143765.141 | 59297.875 | 2.424 | 1909.3 | -36.4 | 239.679 |
| 4000 ng | 4000 | 3.71 | 146956.5 | 47366.727 | 3.103 | 3325 | -16.9 | 190.533 |
| blank | | 3.58 | 1007.281 | 3169.399 | 0.318 | 180.7 | | 6.299 |
| blank | | 3.58 | 1104.284 | 2604.694 | 0.424 | 246.7 | | 4.869 |
| QC-1 | 750 | 3.71 | 110870.07 | 273212.469 | 0.406 | 235.3 | -68.6 | 771.267 |
| QC-2 | 1500 | 3.71 | 129095.914 | 214402.469 | 0.602 | 360.3 | -76 | 363.354 |
| QC-3 | 2500 | 3.71 | 131100 | 138335 | 0.948 | 592.8 | -76.3 | 240.213 |
| blank | | 3.58 | 1044.252 | 3600.424 | 0.29 | 163.7 | | 4.573 |
| blank | | 3.58 | 1265.189 | 3254.976 | 0.389 | 224.6 | | 4.062 |
| blank | | 3.58 | 1376.996 | 2818.832 | 0.488 | 287.4 | | 4.605 |
| QC-2 | 1500 | 3.71 | 123858.117 | 205209 | 0.604 | 361.3 | -75.9 | 426.279 |
| blank | | 3.59 | 1528.398 | 3074.438 | 0.497 | 292.9 | | 5.034 |
| blank | | 3.59 | 1349.482 | 2633.729 | 0.512 | 302.6 | | 4.1 |
| LNG_P1<br>(unrelated pop. control) | | 3.64 | 8578.104 | 335556.094 | 0.026 | 5.4 | | 20.266 |
| LNG_P2<br>(unrelated pop. control) | | 3.63 | 7032.546 | 334425.938 | 0.021 | 2.7 | | 24.727 |
| 2013-303<br>(Mother - unaffected) | | 3.63 | 7301.782 | 340491.313 | 0.021 | 3 | | 27.306 |
| 2013-302<br>(Proband) | | 3.63 | 5838.648 | 354765.781 | 0.016 | 0 | | 21.033 |
| 2013-307<br>(Brother - unaffected) | | 3.63 | 7407.217 | 343583.031 | 0.022 | 3 | | 16.738 |
| blank | | 3.59 | 1528.398 | 3074.438 | 0.497 | 292.9 | | 5.034 |
| blank | | 3.59 | 1349.482 | 2633.729 | 0.512 | 302.6 | | 4.1 |

**Compound 5: C18- Sphingosine**

| Sample | Std. Conc | RT | Area | IS Area | Response | Conc. | %Dev | S/N |
|---|---|---|---|---|---|---|---|---|
| blank | | 3.47 | 5997.581 | 852.739 | 7.033 | 778.8 | | 20.097 |
| blank | | 3.47 | | 331.18 | 8.162 | 903.8 | | 7.196 |
| Solvent blank | | 3.48 | 2044.057 | 409514.375 | 0.005 | 0.5 | | 5.66 |
| 1 ng | 1 | 3.66 | 1448.667 | 172473.375 | 0.008 | 0.8 | -15.5 | 6.164 |
| 10 ng | 10 | 3.66 | 7768.076 | 167206.703 | 0.046 | 5.1 | -49.4 | 43.601 |
| 25 ng | 25 | 3.66 | 41153.301 | 167522.859 | 0.246 | 27.1 | 8.5 | 192.344 |
| 100 ng | 100 | 3.66 | 147997.766 | 142090.656 | 1.042 | 115.3 | 15.3 | 558.526 |
| 500 ng | 500 | 3.66 | 348268.813 | 79152.766 | 4.4 | 487.1 | -2.6 | 1550.477 |
| 1000 ng | 1000 | 3.66 | 412812.656 | 48456.828 | 8.519 | 943.3 | -5.7 | 1671.349 |
| 2000 ng | 2000 | 3.67 | 390926.031 | 22846.123 | 17.111 | 1894.7 | -5.3 | 1670.699 |
| 3000 ng | 3000 | 3.67 | 462594.844 | 16222.605 | 28.515 | 3157.6 | 5.3 | 1769.22 |
| 4000 ng | 4000 | 3.66 | 291693.781 | 15994.171 | 18.238 | 2019.5 | -49.5 | 1056.332 |
| blank | | 3.48 | 2193.423 | 2001.785 | 1.096 | 121.3 | | 6.517 |
| blank | | 3.48 | 2254.033 | 1243.673 | 1.812 | 200.6 | | 5.609 |
| QC-1 | 750 | 3.67 | 441790.688 | 107433.516 | 4.112 | 455.3 | -39.3 | 1728.806 |
| QC-2 | 1500 | 3.67 | 476562.969 | 47096.883 | 10.119 | 1120.4 | -25.3 | 2366.459 |
| QC-3 | 2500 | 3.67 | 458497.5 | 33673.996 | 13.616 | 1507.7 | -39.7 | 1236.162 |
| blank | | 3.49 | 2241.904 | 1757.163 | 1.276 | 141.2 | | 4.309 |
| blank | | 3.49 | 2833.845 | | | | | 7.809 |
| blank | | 3.49 | 1586.056 | 269.331 | 5.889 | 652 | | 6.947 |
| QC-2 | 1500 | 3.67 | 487893.406 | 75788.789 | 6.438 | 712.8 | -52.5 | 1511.461 |
| blank | | 3.49 | 2742.826 | 63.608 | 43.121 | 4774.9 | | 7.563 |
| blank | | 3.49 | 3058.466 | 251.93 | 12.14 | 1344.3 | | 9.722 |
| LNG_P1 (unrelated pop. control) | | 3.67 | 1840.407 | 237122.609 | 0.008 | 0.8 | | 6.303 |
| LNG_P2 (unrelated pop. control) | | 3.67 | 1240.409 | 257406.516 | 0.005 | 0.4 | | 4.939 |
| 2013-303 (Mother - unaffected) | | 3.67 | 1251.613 | 259903.375 | 0.005 | 0.4 | | 4.046 |
| 2013-302 (Proband) | | 3.67 | 1392.526 | 261104.844 | 0.005 | 0.5 | | 6.494 |
| 2013-307 (Brother - unaffected) | | 3.66 | 1393.62 | 251903.094 | 0.006 | 0.5 | | 4.666 |
| blank | | 3.49 | 2742.826 | 63.608 | 43.121 | 4774.9 | | 7.563 |
| blank | | 3.49 | 3058.466 | 251.93 | 12.14 | 1344.3 | | 9.722 |

**Compound 6: C18- Sphinganine**

| Sample | Std. Conc | RT | Area | IS Area | Response | Conc. | %Dev | S/N |
|---|---|---|---|---|---|---|---|---|
| blank | | 3.67 | 7365.816 | | | | | 85.109 |
| blank | | 3.67 | 2067.938 | 66.267 | 31.206 | 573.6 | | 17.864 |
| Solvent blank | | 3.67 | 127661.414 | 238672.281 | 0.535 | 10 | | 1181.289 |
| 1 ng | 1 | 3.67 | 5417.912 | 120841.625 | 0.045 | 1 | 4.4 | 151.266 |
| 10 ng | 10 | 3.67 | 18604.463 | 119027.18 | 0.156 | 3.1 | -69.1 | 332.243 |
| 25 ng | 25 | 3.68 | 123998.789 | 114278.586 | 1.085 | 20.2 | -19.4 | 1218.627 |
| 100 ng | 100 | 3.67 | 424884 | 73346.258 | 5.793 | 106.7 | 6.7 | 1458.093 |
| 500 ng | 500 | 3.67 | 1092794.875 | 34719.465 | 31.475 | 578.6 | 15.7 | 677.842 |
| 1000 ng | 1000 | 3.68 | 1388979.75 | 26643.355 | 52.132 | 958.1 | -4.2 | 389.356 |
| 2000 ng | 2000 | 3.68 | 1607158.375 | 15480.133 | 103.821 | 1907.9 | -4.6 | 232.338 |
| 3000 ng | 3000 | 3.68 | 1731935.25 | 12034.784 | 143.911 | 2644.6 | -11.8 | 145.268 |
| 4000 ng | 4000 | 3.68 | 1730683.75 | 7845.722 | 220.589 | 4053.5 | 1.3 | 128.704 |
| blank | | 3.68 | 26188.396 | 316.611 | 82.715 | 1520.1 | | 229.507 |
| blank | | 3.68 | 8225.282 | 101.716 | 80.865 | 1486.1 | | 93.679 |
| QC-1 | 750 | 3.68 | 1028187.375 | 71626.445 | 14.355 | 264 | -64.8 | 336.789 |
| QC-2 | 1500 | 3.68 | 1306212.25 | 47314.863 | 27.607 | 507.5 | -66.2 | 174.781 |
| QC-3 | 2500 | 3.68 | 1683976 | 28141.639 | 59.839 | 1099.8 | -56 | 161.701 |
| blank | | 3.68 | 18353.52 | 333.439 | 55.043 | 1011.6 | | 150.72 |
| blank | | 3.68 | 7034.597 | 202.57 | 34.727 | 638.3 | | 52.953 |
| blank | | 3.68 | 2337.303 | 65.332 | 35.776 | 657.6 | | 27.732 |
| QC-2 | 1500 | 3.68 | 1289915.5 | 45919.152 | 28.091 | 516.4 | -65.6 | 184.976 |
| blank | | 3.69 | 2225.452 | 42.084 | 52.881 | 971.9 | | 29.478 |
| blank | | 3.69 | 2403.859 | 526.007 | 4.57 | 84.2 | | 24.346 |
| LNG_P1 (unrelated pop. control) | | 3.69 | 4648.483 | 132944.875 | 0.035 | 0.9 | | 55.302 |
| LNG_P2 (unrelated pop. control) | | 3.68 | 2979.833 | 128008.297 | 0.023 | 0.6 | | 37.969 |
| 2013-303 (Mother - unaffected) | | 3.68 | 2982.664 | 131258.594 | 0.023 | 0.6 | | 32.594 |
| 2013-302 (Proband) | | 3.68 | 3650.995 | 141221.547 | 0.026 | 0.7 | | 78.143 |
| 2013-307 (Brother - unaffected) | | 3.68 | 3243.934 | 138557.219 | 0.023 | 0.7 | | 46.535 |
| blank | | 3.69 | 2225.452 | 42.084 | 52.881 | 971.9 | | 29.478 |
| blank | | 3.69 | 2403.859 | 526.007 | 4.57 | 84.2 | | 24.346 |

**Compound 7: C20 Sphingosine**

| Sample | Std. Conc | RT | Area | IS Area | Response | Conc. | %Dev | S/N |
|---|---|---|---|---|---|---|---|---|
| blank | | 3.71 | 2304.052 | 852.739 | 2.702 | 347.2 | | 12.777 |
| blank | | 3.71 | 306.437 | 331.18 | 0.925 | 118.8 | | 2.916 |
| Solvent blank | | 3.71 | 41232.457 | 409514.375 | 0.101 | 12.8 | | 475.476 |
| 1 ng | 1 | 3.71 | 1557.2 | 172473.375 | 0.009 | 1.1 | 5.2 | 25.073 |
| 10 ng | 10 | 3.71 | 5509.329 | 167206.703 | 0.033 | 4.1 | -58.7 | 44.947 |
| 25 ng | 25 | 3.72 | 34747.246 | 167522.859 | 0.207 | 26.6 | 6.2 | 369.48 |
| 100 ng | 100 | 3.72 | 106314.977 | 142090.656 | 0.748 | 96.1 | -3.9 | 735.755 |
| 500 ng | 500 | 3.72 | 283312.219 | 79152.766 | 3.579 | 460 | -8 | 4174.237 |
| 1000 ng | 1000 | 3.72 | 364705.469 | 48456.828 | 7.526 | 967.4 | -3.3 | 3032.124 |
| 2000 ng | 2000 | 3.72 | 368816.625 | 22846.123 | 16.144 | 2075.1 | 3.8 | 3352.16 |
| 3000 ng | 3000 | 3.73 | 378587.281 | 16222.605 | 23.337 | 2999.8 | 0 | 2966.696 |
| 4000 ng | 4000 | 3.73 | 394485.063 | 15994.171 | 24.664 | 3170.4 | -20.7 | 2918.403 |
| blank | | 3.72 | 13875.337 | 2001.785 | 6.931 | 890.9 | | 133.762 |
| blank | | 3.72 | 4447.397 | 1243.673 | 3.576 | 459.6 | | 34.703 |
| QC-1 | 750 | 3.72 | 320821.313 | 107433.516 | 2.986 | 383.8 | -48.8 | 2910.874 |
| QC-2 | 1500 | 3.73 | 387077.469 | 47096.883 | 8.219 | 1056.4 | -29.6 | 5252.003 |
| QC-3 | 2500 | 3.73 | 391747.75 | 33673.996 | 11.634 | 1495.4 | -40.2 | 2781.323 |
| blank | | 3.72 | 9708.028 | 1757.163 | 5.525 | 710.1 | | 81.365 |
| blank | | 3.72 | 3231.016 | | | | | 32.962 |
| blank | | 3.72 | 532.438 | 269.331 | 1.977 | 254 | | 3.811 |
| QC-2 | 1500 | 3.73 | 382505.781 | 75788.789 | 5.047 | 648.7 | -56.8 | 3546.115 |
| blank | | 3.72 | 594.587 | 63.608 | 9.348 | 1201.5 | | 5.873 |
| blank | | 3.72 | 537.572 | 251.93 | 2.134 | 274.2 | | 4.86 |
| LNG_P1 (unrelated pop. control) | | 3.72 | 1705.062 | 237122.609 | 0.007 | 0.8 | | 12.409 |
| LNG_P2 (unrelated pop. control) | | 3.72 | 999.18 | 257406.516 | 0.004 | 0.4 | | 4.908 |
| 2013-303 (Mother - unaffected) | | 3.72 | 1063.219 | 259903.375 | 0.004 | 0.4 | | 3.751 |
| 2013-302 (Proband) | | 3.72 | 1067.746 | 261104.844 | 0.004 | 0.4 | | 5.06 |
| 2013-307 (Brother - unaffected) | | 3.72 | 680.237 | 251903.094 | 0.003 | 0.2 | | 4.577 |
| blank | | 3.72 | 594.587 | 63.608 | 9.348 | 1201.5 | | 5.873 |
| blank | | 3.72 | 537.572 | 251.93 | 2.134 | 274.2 | | 4.86 |

**Compound 8: C20 Sphinganine**

| Sample | Std. Conc | RT | Area | IS Area | Response | Conc. | %Dev | S/N |
|---|---|---|---|---|---|---|---|---|
| blank | | 3.72 | 5623.247 | | | | | 77.99 |
| blank | | 3.72 | 1460.552 | 66.267 | 22.04 | 328.1 | | 28.868 |
| Solvent blank | | 3.72 | 101370.625 | 238672.281 | 0.425 | 7.1 | | 1596.882 |
| 1 ng | 1 | 3.72 | 3183.616 | 120841.625 | 0.026 | 1.2 | 18.7 | 134.039 |
| 10 ng | 10 | 3.72 | 9156.259 | 119027.18 | 0.077 | 1.9 | -80.6 | 217.392 |
| 25 ng | 25 | 3.72 | 69727.141 | 114278.586 | 0.61 | 9.9 | -60.6 | 1366.242 |
| 100 ng | 100 | 3.72 | 246611.859 | 73346.258 | 3.362 | 50.7 | -49.3 | 3810.613 |
| 500 ng | 500 | 3.72 | 984110.5 | 34719.465 | 28.345 | 421.8 | -15.6 | 1535.163 |
| 1000 ng | 1000 | 3.73 | 1622324 | 26643.355 | 60.89 | 905.1 | -9.5 | 1308.659 |
| 2000 ng | 2000 | 3.73 | 2122365 | 15480.133 | 137.103 | 2037.1 | 1.9 | 986.358 |
| 3000 ng | 3000 | 3.73 | 2540340.25 | 12034.784 | 211.083 | 3135.8 | 4.5 | 863.315 |
| 4000 ng | 4000 | 3.73 | 2730303.5 | 7845.722 | 347.999 | 5169.3 | 29.2 | 749.734 |
| blank | | 3.72 | 31931.826 | 316.611 | 100.855 | 1498.7 | | 325.994 |
| blank | | 3.72 | 8966.357 | 101.716 | 88.151 | 1310 | | 118.892 |
| QC-1 | 750 | 3.73 | 1366073.25 | 71626.445 | 19.072 | 284.1 | -62.1 | 1763.017 |
| QC-2 | 1500 | 3.73 | 2324283.25 | 47314.863 | 49.124 | 730.4 | -51.3 | 1027.326 |
| QC-3 | 2500 | 3.73 | 2647881.5 | 28141.639 | 94.091 | 1398.3 | -44.1 | 942.374 |
| blank | | 3.73 | 23491.076 | 333.439 | 70.451 | 1047.1 | | 167.398 |
| blank | | 3.73 | 7284.76 | 202.57 | 35.962 | 534.9 | | 152.583 |
| blank | | 3.73 | 1422 | 65.332 | 21.766 | 324.1 | | 25.123 |
| QC-2 | 1500 | 3.73 | 2250878.25 | 45919.152 | 49.018 | 728.8 | -51.4 | 1152.206 |
| blank | | 3.73 | 1245.154 | 42.084 | 29.587 | 440.2 | | 35.427 |
| blank | | 3.73 | 1312.826 | 526.007 | 2.496 | 37.9 | | 46.359 |
| LNG_P1 (unrelated pop. control) | | 3.73 | 3440.664 | 132944.875 | 0.026 | 1.2 | | 79.243 |
| LNG_P2 (unrelated pop. control) | | 3.73 | 2577.307 | 128008.297 | 0.02 | 1.1 | | 59.189 |
| 2013-303 (Mother - unaffected) | | 3.72 | 2799.236 | 131258.594 | 0.021 | 1.1 | | 72.301 |
| 2013-302 (Proband) | | 3.73 | 2746.015 | 141221.547 | 0.019 | 1.1 | | 62.369 |
| 2013-307 (Brother - unaffected) | | 3.72 | 2373.536 | 138557.219 | 0.017 | 1.1 | | 62.376 |
| blank | | 3.73 | 1245.154 | 42.084 | 29.587 | 440.2 | | 35.427 |
| blank | | 3.73 | 1312.826 | 526.007 | 2.496 | 37.9 | | 46.359 |

**Compound 9: C17 Sphingosine**

| Sample | Std. Conc | RT | Area | IS Area | Response | Conc. | %Dev | S/N |
|---|---|---|---|---|---|---|---|---|
| blank | | 3.63 | 1064.061 | 852.739 | 1.248 | 78.5 | | 2.416 |
| blank | | 3.63 | 271.214 | 331.18 | 0.819 | 51.5 | | 0.499 |
| Solvent blank | | 3.64 | 57684.277 | 409514.375 | 0.141 | 9.1 | | 171.491 |
| 1 ng | 1 | 3.64 | 2336.133 | 172473.375 | 0.014 | 1.2 | 24.7 | 8.162 |
| 10 ng | 10 | 3.64 | 11219.798 | 167206.703 | 0.067 | 4.6 | -54.3 | 24.957 |
| 25 ng | 25 | 3.64 | 58254.777 | 167522.859 | 0.348 | 22 | -11.9 | 150.324 |
| 100 ng | 100 | 3.64 | 195291.313 | 142090.656 | 1.374 | 86.5 | -13.5 | 530.559 |
| 500 ng | 500 | 3.64 | 576614 | 79152.766 | 7.285 | 479.8 | -4 | 1451.26 |
| 1000 ng | 1000 | 3.64 | 719283.75 | 48456.828 | 14.844 | 1055.3 | 5.5 | 1636.363 |
| 2000 ng | 2000 | 3.64 | 646635.188 | 22846.123 | 28.304 | 2530 | 26.5 | 1694.959 |
| 3000 ng | 3000 | 3.65 | 443568.875 | 16222.605 | 27.343 | 2382.9 | -20.6 | 1141.363 |
| 4000 ng | 4000 | 3.64 | 529962.438 | 15994.171 | 33.135 | 3843.5 | -3.9 | 1884.656 |
| blank | | 3.64 | 5612.765 | 2001.785 | 2.804 | 178 | | 13.582 |
| blank | | 3.64 | 1907.333 | 1243.673 | 1.534 | 96.6 | | 4.815 |
| QC-1 | 750 | 3.64 | 639952.313 | 107433.516 | 5.957 | 387.8 | -48.3 | 1942.57 |
| QC-2 | 1500 | 3.64 | 669295.5 | 47096.883 | 14.211 | 1003.1 | -33.1 | 1676.127 |
| QC-3 | 2500 | 3.65 | 465914.656 | 33673.996 | 13.836 | 972.5 | -61.1 | 1112.196 |
| blank | | 3.64 | 3673.259 | 1757.163 | 2.09 | 132.1 | | 9.065 |
| blank | | 3.64 | 1331.779 | | | | | 3.222 |
| blank | | 3.64 | 615.738 | 269.331 | 2.286 | 144.6 | | 1.056 |
| QC-2 | 1500 | 3.64 | 688140.438 | 75788.789 | 9.08 | 607.9 | -59.5 | 1623.612 |
| blank | | 3.64 | 374.6 | 63.608 | 5.889 | 383.2 | | 0.86 |
| blank | | 3.64 | 274.25 | 251.93 | 1.089 | 68.4 | | 0.734 |
| LNG_P1<br>(unrelated pop. control) | | 3.65 | 1225.391 | 237122.609 | 0.005 | 0.7 | | 2.023 |
| LNG_P2<br>(unrelated pop. control) | | 3.64 | 1129.99 | 257406.516 | 0.004 | 0.7 | | 1.354 |
| 2013-303<br>(Mother - unaffected) | | 3.64 | 1133.067 | 259903.375 | 0.004 | 0.7 | | 1.653 |
| 2013-302<br>(Proband) | | 3.64 | 964.462 | 261104.844 | 0.004 | 0.6 | | 1.4 |
| 2013-307<br>(Brother - unaffected) | | 3.64 | 1230.287 | 251903.094 | 0.005 | 0.7 | | 1.197 |
| blank | | 3.64 | 374.6 | 63.608 | 5.889 | 383.2 | | 0.86 |
| blank | | 3.64 | 274.25 | 251.93 | 1.089 | 68.4 | | 0.734 |

**Table S3. *SPTLC1* mutations identified in ALS cases**

| SNP information | | No. cases | Population frequency | | Prediction and conservation algorithms | | |
|---|---|---|---|---|---|---|---|
| | | | gnomAD | HRC | Prediction | Conservation | dbscSNV |
| 9:94843196C>T | p.Ala104Thr | 1 | $1{\cdot}1\times10^{-5}$ | 0 | damaging | conserved | NA |
| 9:94842347C>G | p.Lys126Asn | 1 | 0 | 0 | damaging | not conserved | NA |
| 9:94821543C>T | p.Arg203His | 1 | $1{\cdot}1\times10^{-5}$ | 0 | damaging | conserved | NA |
| 9:94821496G>A | p.Arg219X | 1 | $2{\cdot}2\times10^{-5}$ | 0 | damaging | conserved | NA |
| 9:94821459A>G | c.690+2T>C | 1 | 0 | 0 | NA | conserved | 0·994 |
| 9:94817761G>A | p.Arg236Cys | 1 | 0 | 0 | damaging | not conserved | NA |
| 9:94817749G>A | p.Arg240Cys | 1 | $1{\cdot}1\times10^{-5}$ | 0 | damaging | conserved | NA |
| 9:94809966C>T | p.Ala305Thr | 3 | $1{\cdot}1\times10^{-5}$ | 0 | damaging | conserved | NA |
| 9:94809927A>T | p.Cys318Ser | 1 | $1{\cdot}1\times10^{-5}$ | 0 | damaging | conserved | NA |
| 9:94809550G>A | p.Arg329X | 1 | $1{\cdot}1\times10^{-5}$ | 0 | damaging | conserved | 0·15 |
| 9:94800603A>G | p.Phe394Ser | 1 | 0 | 0 | damaging | conserved | NA |
| 9:94800576C>T | p.Gly403Glu | 1 | 0 | 0 | damaging | conserved | NA |
| 9:94800568C>T | p.Glu406Lys | 1 | 0 | 0 | damaging | conserved | NA |
| 9:94797161A>G | p.Met420Thr | 1 | 0 | 0 | damaging | conserved | NA |
| 9:94794835C>T | p.Arg445Gln | 1 | $2{\cdot}2\times10^{-5}$ | 0 | damaging | conserved | NA |
| 9:94794766G>A | p.Ala468Val | 1 | $1{\cdot}1\times10^{-5}$ | 0 | damaging | conserved | NA |
| 9:94794610A>- | p.Tyr509X | 1 | 0 | 0 | damaging | conserved | NA |

Variant position is in build 19; amino acid change is based on the canonical transcript NP_006406, except for p.Tyr509X which is based on NP_001268232; No. cases, number of ALS cases carrying the variant; only variants with frequency less than $3{\cdot}3\times10^{-5}$ are shown; gnomAD, frequency shown is the maximum in either the Finnish or the non-Finnish European populations (non-neurological samples, version 2·1); HRC, Haplotype Reference Consortium (version 1); 'damaging' means the variant is designated as such by at least four out of five prediction algorithms consisting of FATHMM, M-CAP, MetaLR, MetaSVM and VEST3[5]; 'conserved' means the position is considered as such by the GERP++, phastCons100way_vertebrate, phyloP100way_vertebrate, and SiPhy_29way algorithms; dbscSNV adaptive boosting score greater than 0·6 indicates a deleterious splice mutation; NA, not available.

**Table S4. Primer sequences used for Sanger sequencing of *SPTLC1* (NM_006415)**

PCR and sequencing primers

| | |
|---|---|
| 1 forward | GGTCTCAGGCCACAAATCC |
| 1 reverse | CACCTGACCCCGTCCAG |
| 2 forward | GAGCCACCACAAATCCTTTC |
| 2 reverse | AAATTTTATGTGCAGGTGTTAGAAG |
| 3 forward | CTCCTGGGAACTTCAAAATG |
| 3 reverse | GAGTGACAGCGAACTTGAGAATAG |
| 4 forward | CCCAGCCTGCCTTTGTC |
| 4 reverse | TTGTTGTGCCTACCACCAAG |
| 5 forward | GCCCAGCCTAAAATTGAATC |
| 5 reverse | TGGGTTAAAGAATAGGAAATGTTTG |
| 6 forward | TGAAACCTTTGGTGAGGTG |
| 6 reverse | TGGCCATAAGGTGAAAGTATTG |
| 7 forward | AGACTTGGCTCTCCAACACC |
| 7 reverse | CATGGATGAAGAGCAAACAATG |
| 8 forward | CGGTCATAAACAAAAGGCAG |
| 8 reverse | AAGTGCTGGGGAATATGTAACAG |
| 9 forward | GGCCTAGCAGAATGGAACTAC |
| 9 reverse | GAATTCACAAGTATTGAGAGTCAGC |
| 10 forward | GAGGTGACCAATGGAGAAATTAG |
| 10 reverse | TCTCTAGGCTTGAAGTTGGC |
| 11 forward | TGGCAAATAATTTCTCCAGTAAG |
| 11 reverse | TTGTTCTAGAGAACTGGTGCC |
| 12 forward | CTGGTCAAACTGACCCAAGC |
| 12 reverse | AAATTGATGAACTTGATCTGGG |
| 13 forward | AAATTGCTTGGGGCAAGG |
| 13 reverse | GAGCGATCTTCAAGCCTGG |
| 14 forward | TTCAAAAGATAACGTATTTCATAGAGC |
| 14 reverse | GTTGCAGGATTTCAGCGTTC |
| 15 forward | TGAAGCTCCAGTTTTAAACGAC |
| 15 reverse | TGACCTCGTGATCCACCTG |

**Table S5. The haplotype of chromosome 9q22.31 associated with the p.Ala305Thr variant in *SPTLC1***

| | | | | UK ALS0039 | | US ND10247 | | Australia AUS145-010335 | |
|---|---|---|---|---|---|---|---|---|---|
| **Marker** | **Position** | **Gene** | **MAF** | **Major** | **Minor** | **Major** | **Minor** | **Major** | **Minor** |
| rs7019926 | 94118106 | *AUH* | 0.24 | T | T | T | **C** | T | T |
| rs10992065 | 94487459 | *ROR2* | 0.26 | C | **C** | C | **T** | C | **T** |
| rs10820900 | 94495608 | *ROR2* | 0.38 | C | **C** | T | **C** | C | **C** |
| rs7855522 | 94499618 | *ROR2* | 0.44 | A | **A** | G | **A** | A | **A** |
| rs12683181 | 94518328 | *ROR2* | 0.33 | T | **T** | C | **T** | T | **T** |
| p.A305T | 94809966 | *SPTLC1* | NA | C | **T** | C | **T** | C | **T** |
| rs7858880 | 94919672 | *LINC00475* | 0.10 | G | G | G | **C** | G | G |

The three SPTLC1 p.Ala305Thr carriers carry a maximal common haplotype spanning 43kb, suggesting a possible ancestral founder. The minimal region is defined by SNP markers rs10820900, rs7855522, and rs12683181 that also includes the p.Ala305Thr mutant T allele (C-A-T-T).

### Other supplementary files

**Movie S1. Tongue fasciculations and wasting in patient 2**
**Movie S2. Gower's sign in patient 2**
**Movie S3. Neurological manifestations in patient 3**
**Movie S4. MRI brain and spinal cord in patient 2**

**Consortia authors and affiliations**

**The members of the ITALSGEN Consortium are:**
Francesco Pio Ausiello[1], Marco Barberis[2], Ilaria Bartolomei[3], Stefania Battistini[4], Michele Benigni[4], Enrica Bersano[5], Giuseppe Borghero[6], Maura Brunetti[7], Andrea Calvo[2], Antonino Cannas[6], Antonio Canosa[2], Margherita Capasso[8], Claudia Caponnetto[9], Patrizio Cardinali[10], Paola Carrera[11], Federico Casale[2], Adriano Chiò[2], Tiziana Colletti[12], Francesca L Conforti[13], Amelia Conte[14], Elisa Conti[15], Massimo Corbo[16], Eleonora Dalla Bella[5], Giovanni Defazio[6], Raffaele Dubbioso[1], Antonio Fasano[17], Cinzia Femiano[18], Carlo Ferrarese[15], Nicola Fini[17], Gianluca Floris[6], Giuseppe Fuda[2], Fabio Giannini[4], Carlo Guidi[19], Antonio Ilardi[2], Vincenzo La Bella[12], Serena Lattante[14], Giuseppe Lauria[5,20], Giancarlo Logroscino[21], Francesco O Logullo[22], Christian Lunetta[23], Gianluigi Mancardi[9], Paola Mandich[9], Jessica Mandrioli[17], Umberto Manera[2], Giuseppe Marangi[14], Kalliopi Marinou[24], Maria Giovanna Marrosu[6], Maurizio Melis[6], Sonia Messina[25], Cristina Moglia[2], Maria Rosaria Monsurrò[18], Gabriele Mora[26], Lorena Mosca[27], Maria Rita Murru[6], Patrizia Occhineri[28], Paola Origone[9], Antonio Petrucci[29], Giovanni Piccirillo[18], Angelo Pirisi[28], Maura Pugliatti[28], Gabriella Restagno[7], Claudia Ricci[4], Nilo Riva[11], Massimo Russo[30], Mario Sabatelli[14], Gessica Sala[15], Fabrizio Salvi[3], Marialuisa Santarelli[31], Lucio Santoro[1], Riccardo Sideri[24], Isabella Simone[21], Rossella Spataro[12], Raffaella Tanel[32], Gioacchino Tedeschi[18], Anna Ticca[33], Antonella Torriello[34], Maria Claudia Torrieri[2], Lucio Tremolizzo[15], Francesca Trojsi[18], Rosario Vasta[2], Giuseppe Vita[25], Paolo Volanti[35], Marcella Zollino[14,20]

1. Università degli Studi di Napoli Federico II, Naples, Italy.
2. Rita Levi Montalcini' Department of Neuroscience, Amyotrophic Lateral Sclerosis Center, University of Turin, Turin, Italy.
3. Center for Diagnosis and Cure of Rare Diseases, Department of Neurology, IRCCS Institute of Neurological Sciences, Bologna, Italy.
4. Department of Medical, Surgical and Neurological Sciences, University of Siena, Siena, Italy.
5. Neuroalgology and Headache Unit, Fondazione IRCCS Istituto Neurologico "Carlo Besta", Milan, Italy.
6. Department of Neurology, Azienda Universitario Ospedaliera di Cagliari and University of Cagliari, Cagliari, Italy.
7. Molecular Genetics Unit, Department of Clinical Pathology, A.S.O. O.I.R.M.-S. Anna, 10126 Turin, Italy
8. Department of Neurology, University of Chieti, Chieti, Italy.
9. Department of Neurosciences, Ophthalmology, Genetics, Rehabilitation, Maternal and Child Health, IRCCS Azienda Ospedaliero-Universitaria San Martino IST, Genoa, Italy.
10. Amministrazione ASUR Zona Territoriale 11, Fermo, Italy.
11. Department of Neurology and Institute of Experimental Neurology (INSPE), IRCCS San Raffaele Scientific Institute, Milan, Italy.
12. ALS Clinical Research Center, Bio. Ne. C., University of Palermo, Palermo, Italy.
13. Institute of Neurological Sciences, National Research Council, Mangone, Cosenza, Italy.
14. Centro Clinico NEMO-Roma, Neurological Institute, Catholic University and I.C.O.M.M. Association for ALS Research, Rome, Italy.
15. Neurology Unit, School of Medicine and Surgery and NeuroMI, University of Milano-Bicocca, Monza, Italy.
16. Department of Neurorehabilitation Sciences (P.T., M.C.), Casa Cura Policlinico, Milan, Italy.
17. Department of Neuroscience, S. Agostino-Estense Hospital, University of Modena and Reggio Emilia, Modena, Italy.
18. Department of Medical, Surgical Neurological Metabolic and Aging Sciences, Second University of Naples, Naples, Italy.
19. AUSL della Romagna, Forlì, Italy.
20. Department of Biomedical and Clinical Sciences "Luigi Sacco", University of Milan, Milan, Italy.
21. Department of Basic Medical Sciences, Neurosciences and Sense Organs, University of Bari, Bari, Italy.
22. Neurological Clinic, Marche Polytechnic University, Ancona, Italy.
23. NeuroMuscular Omnicenter, Serena Onlus Foundation, Milan, Italy.
24. Department of Neurological Rehabilitation, Fondazione Salvatore Maugeri, IRCCS, Istituto Scientifico di Milano, Milan, Italy.
25. OUC Neurology and Neuromuscular Disorders, University of Messina, Italy.
26. ALS Center, ICS Maugeri, IRCCS, Milan, Italy.
27. Department of Laboratory Medicine, Medical Genetics, Niguarda Ca' Granda Hospital, Milan, Italy.
28. Department of Biomedical and Surgical Sciences, Section of Neurological, Psychiatric and Psychological Sciences, University of Ferrara, Ferrara, Italy.
29. Neurology Department, San Camillo Hospital, Rome, Italy.
30. Centro Clinico NEMO-Messina, Messina, Italy.
31. Department of Medicine, Azienda Complesso Ospedaliero, San Filippo Neri, Rome, Italy.
32. Department of Neurology, Santa Chiara Hospital, Trento, Italy.
33. Department of Neurology, Azienda Ospedaliera San Francesco, Nuoro, Italy.

34. AOU OO.RR. San Giovanni di Dio Ruggi d'Aragona Salerno, Salerno, Italy.
35. Neurorehabilitation Unit/ALS Center, Scientific Clinical Institutes (ICS) Maugeri, IRCCS, Mistretta, Messina, Italy.

**The members of the international ALS genomics consortium (iALSgc) are:**

Yevgeniya Abramzon[1,2], Sampath Arepalli[3], Robert H. Baloh[4], Robert Bowser[5], Christopher B. Brady[6], Alexis Brice[7,8], James Broach[9], Roy H. Campbell[10], William Camu[11], Ruth Chia[1], Adriano Chiò[12,13], John Cooper-Knock[14], Jinhui Ding[15], Carsten Drepper[16], Vivian E. Drory[17], Travis L. Dunckley[18], John D. Eicher[19], Faraz Faghri[20,10], Eva Feldman[21], Mary Kay Floeter[22], Pietro Fratta[2], Joshua T. Geiger[23], Glenn Gerhard[18], J. Raphael Gibbs[15], Summer B. Gibson[24], Jonathan D. Glass[25], John Hardy[26], Matthew B. Harms[27], Terry D. Heiman-Patterson[28,29], Dena G. Hernandez[3], Lilja Jansson[30], Freya Kamel[31], Janine Kirby[14], Neil W. Kowall[32], Hannu Laaksovirta[30], John E. Landers[33], Francesco Landi[34], Isabelle Le Ber[7,8], Serge Lumbroso[35], Daniel JL. MacGowan[36], Nicholas J. Maragakis[37], Gabriele Mora[38], Kevin Mouzat[35], Natalie A. Murphy[1], Liisa Myllykangas[39], Mike A. Nalls[20,40], Richard W. Orrell[41], Lyle W. Ostrow[37], Roger Pamphlett[42], Stuart Pickering-Brown[43], Erik P. Pioro[44], Hannah A. Pliner[1], Stefan M. Pulst[24], John M. Ravits[45], Alan E. Renton[1,46], Alberto Rivera[1], Wim Robberecht[47], Ekaterina Rogaeva[48], Sara Rollinson[43], Jeffrey D. Rothstein[37], Sonja W. Scholz[23,37], Michael Sendtner[49], Pamela J. Shaw[14], Katie C. Sidle[26], Zachary Simmons[50], Andrew B. Singleton[20], David J. Stone[19], Pentti J. Tienari[30], Bryan J. Traynor[1,37], John Q. Trojanowski[51], Juan C. Troncoso[52], Miko Valori[30], Philip Van Damme[47,53], Vivianna M. Van Deerlin[51], Ludo Van Den Bosch[47], Lorne Zinman[54]

1. Neuromuscular Diseases Research Section, Laboratory of Neurogenetics, National Institute on Aging, Bethesda, MD, 20892, USA.
2. Sobell Department of Motor Neuroscience and Movement Disorders, Institute of Neurology, University College London, London, WC1N 3BG, UK.
3. Genomics Technology Group, Laboratory of Neurogenetics, National Institute on Aging, Bethesda, MD, 20892, USA.
4. Department of Neurology, Cedars-Sinai Medical Center, Los Angeles, CA, 90048, USA.
5. Division of Neurology, Barrow Neurological Institute, Phoenix, AZ, 85013, USA.
6. Research and Development Service, Veterans Affairs Boston Healthcare System, Boston, MA, 02130, USA.
7. Centre de Recherche de l'Institut du Cerveau et de la Moelle épinière, Université Pierre et Marie Curie, Paris, France.
8. INSERM U975, Paris, France.
9. Department of Biochemistry, Penn State College of Medicine, Hershey, PA, 17033, USA.
10. Department of Computer Science, University of Illinois at Urbana-Champaign, 201 North Goodwin Avenue, Urbana, IL, 61801, USA.
11. ALS Reference Center, Gui de Chauliac Hospital, CHU and Univ Montpellier, Montpellier France, Montpellier, France.
12. 'Rita Levi Montalcini' Department of Neuroscience, University of Turin, Via Verdi 8, Turin, 10124, Italy.
13. Neuroscience Institute of Torino, University of Turin, Turin, 10124, Italy.
14. Department of Neuroscience, University of Sheffield, Sheffield, S10 2HQ, UK.
15. Computational Biology Core, Laboratory of Neurogenetics, National Institute on Aging, Bethesda, MD, 20892, USA.
16. Institute for Clinical Neurobiology, University of Würzburg, Würzburg, D-97078, Germany.
17. Department of Neurology, Tel-Aviv Sourasky Medical Center, Tel-Aviv, Israel.
18. Department of Pathology, Penn State College of Medicine, Hershey, PA, 17033, USA.
19. Genetics, Genetics and Pharmacogenomics, Merck Research Laboratories, Merck & Co., Inc., West Point, PA, 19486, USA.
20. Molecular Genetics Section, Laboratory of Neurogenetics, National Institute on Aging, Bethesda, MD, 20892, USA.
21. Department of Neurology, University of Michigan, 1500 E Medical Center Dr, Ann Arbor, MI, 48109, USA.
22. Motor Neuron Disorders Unit, Laboratory of Neurogenetics, National Institute of Neurological Disorders and Stroke, Bethesda, MD, 20892, USA.
23. Neurodegenerative Diseases Research Unit, Laboratory of Neurogenetics, National Institute of Neurological Disorders and Stroke, Bethesda, MD, 20892, USA.
24. Department of Neurology, University of Utah School of Medicine, 175 North Medical Drive East, Salt Lake City, UT, 84132, USA.
25. Department of Neurology, Emory University School of Medicine, Atlanta, GA, 30322, USA.
26. Department of Molecular Neuroscience and Reta Lila Weston Laboratories, Institute of Neurology, University College London, London, WC1N 3BG, UK.
27. Department of Neurology, Columbia University, New York, NY, 10032, USA.
28. Department of Neurology, Drexel University College of Medicine, Philadelphia, PA, 19102, USA.

29. Department of Neurology, Temple University, 7602 Central Ave, Philadelphia, PA, 19111, USA.
30. Department of Neurology, University of Helsinki, Helsinki, FIN-02900, Finland.
31. Epidemiology Branch, National Institute of Environmental Health Sciences, Durham, NC, 27709, USA.
32. Department of Neurology, Veterans Affairs Boston Healthcare System, Boston, MA, 02130, USA.
33. Department of Neurology, University of Massachusetts Medical School, Worcester, MA, 01605, USA.
34. Department of Geriatrics, Neurosciences and Orthopedics, Center for Geriatric Medicine, Catholic University of Sacred Heart, Rome, 00168, Italy.
35. Service de Biochimie, CHU de Nîmes, Nîmes, France.
36. Neuromuscular Division and ALS Center, Beth Israel Medical Center, Albert Einstein College of Medicine, New York, NY, 01605, USA.
37. Department of Neurology, Johns Hopkins University, Baltimore, MD, 21287, USA.
38. ALS Center, ICS Maugeri, IRCCS, Via Camaldoli, 64, Milan, 20138, Italy.
39. Department of Pathology, Haartman Institute/HUSLAB, University of Helsinki and Folkhalsan Research Center (LM), Helsinki, FIN-02900, Finland.
40. Data Tecnica International, Glen Echo, MD, 20812, USA.
41. Department of Clinical Neuroscience, Institute of Neurology, University College London, London, NW2 2PG, UK.
42. Discipline of Pathology, Brain and Mind Centre, University of Sydney, Camperdown, NSW 2050, Australia.
43. Faculty of Human and Medical Sciences, University of Manchester, Manchester, M13 9PT, UK.
44. Department of Neurology, Cleveland Clinic, Cleveland, OH, 44195, USA.
45. Department of Neuroscience, Experimental Neurology and Leuven Research Institute for Neuroscience and Disease, University of California San Diego, 9500 Gilman Drive, La Jolla, CA, 92093, USA.
46. Nash Family Department of Neuroscience, Ronald M. Loeb Center for Alzheimer's Disease, Icahn School of Medicine at Mount Sinai, New York, NY, 10029, USA.
47. Department of Neurosciences, Experimental Neurology and Leuven Research Institute for Neuroscience and Disease, University of Leuven, Leuven, 3000, Belgium.
48. Division of Neurology, Tanz Centre for Research of Neurodegenerative Diseases and Toronto Western Hospital, University of Toronto, Toronto, M5S 3H2, Canada.
49. Department of Neurology, Institute for Clinical Neurobiology, University of Würzburg, Würzburg, D-97078, Germany.
50. Department of Neurology, Penn State College of Medicine, Hershey, PA, 17033, USA.
51. Department of Pathology and Laboratory Medicine, University of Pennsylvania, Philadelphia, PA, 19104, USA.
52. Clinical and Neuropathology Core, Johns Hopkins University, Baltimore, MD, 21287, USA.
53. VIB, Center for Brain & Disease Research, Laboratory of Neurobiology, University of Leuven, Leuven, 3000, Belgium.
54. Division of Neurology, Sunnybrook Health Sciences Centre, University of Toronto, Toronto, M4N 3M5, Canada.

**The members of the FALS Sequencing Consortium are:**

Bradley N. Smith[1], Nicola Ticozzi[2,3], Claudia Fallini[4], Athina Soragia Gkazi[1], Simon D. Topp[1,5], Emma L. Scotter[6], Kevin P. Kenna[4], Pamela Keagle[4], Jack W. Miller[7], Cinzia Tiloca[8], Caroline Vance[9], Claire Troakes[9], Claudia Colombrita[8], Safa Al-Sarraj[1], Andrew King[1], Daniela Calini[2], Viviana Pensato[10], Barbara Castellotti[10], Jacqueline de Belleroche[11], Frank Baas[12], Anneloor L.M.A. ten Asbroek[13], Peter C. Sapp[4], Diane McKenna-Yasek[4], Russell L. McLaughlin[14], Meraida Polak[15], Seneshaw Asress[15], Jesús Esteban-Pérez[16], José Luis Muñoz-Blanco[17], Zorica Stevic[18], Sandra D'Alfonso[19], Letizia Mazzini[20], Giacomo P. Comi[21], Roberto Del Bo[21], Mauro Ceroni[22], Stella Gagliardi[23], Giorgia Querin[24], Cinzia Bertolin[24], Wouter van Rheenen[25], Frank P. Diekstra[25], Rosa Rademakers[26], Marka van Blitterswijk[26], Kevin B. Boylan[27], Giuseppe Lauria[28], Stefano Duga[29], Stefania Corti[21], Cristina Cereda[23], Lucia Corrado[19], Gianni Sorarù[24], Kelly L. Williams[30], Garth A. Nicholson[30], Ian P. Blair[30], Claire Leblond-Manry[31], Guy A. Rouleau[32], Orla Hardiman[33], Karen E. Morrison[34], Jan H. Veldink[25], Leonard H. van den Berg[25], Ammar Al-Chalabi[9], Hardev Pall[35], Pamela J. Shaw[36], Martin R. Turner[7], Kevin Talbot[7], Franco Taroni[10], Alberto García-Redondo[16], Zheyang Wu[37], Jonathan D. Glass[15], Cinzia Gellera[10], Antonia Ratti[2], Robert H. Brown, Jr.[4], Vincenzo Silani[2,3], Christopher E. Shaw[9,5], John E. Landers[4]

1. Centre for Neurodegeneration Research, King's College London, Department of Clinical Neuroscience, Institute of Psychiatry, London, UK.
2. Department of Neurology and Laboratory of Neuroscience, IRCCS Istituto Auxologico Italiano, Milan 20149, Italy.
3. Department of Pathophysiology and Transplantation, "Dino Ferrari" Center, Università degli Studi di Milano, Milan 20122, Italy.
4. Department of Neurology, University of Massachusetts Medical School, Worcester, Massachusetts 01605, USA.
5. UK Dementia Research Institute at King's College London, London, UK.

6. Centre for Brain Research, University of Auckland, New Zealand.
7. Nuffield Department of Clinical Neurosciences, University of Oxford, UK.
8. Department of Neurology, IRCCS Istituto Auxologico Italiano, Milan, Italy.
9. Maurice Wohl Clinical Neuroscience Institute, Department of Basic and Clinical Neuroscience, King's College London, London SE5 9RS, UK.
10. Unit of Medical Genetics and Neurogenetics, Fondazione IRCCS Istituto Neurologico 'Carlo Besta', Milan 20133, Italy.
11. Neurogenetics Group, Division of Brain Sciences, Hammersmith Hospital Campus, Burlington Danes Building, Du Cane Road, London, UK.
12. Clinical Genetics, Leiden University Medical Center, Leiden, the Netherlands.
13. Department of Neurogenetics and Neurology, Academic Medical Centre, Amsterdam, the Netherlands.
14. Population Genetics Laboratory, Smurfit Institute of Genetics, Trinity College Dublin, Dublin, Republic of Ireland.
15. Department of Neurology, Emory University, Atlanta, GA 30322, USA.
16. Unidad de ELA, Instituto de Investigación Hospital 12 de Octubre de Madrid, SERMAS, and Centro de Investigación Biomédica en Red de Enfermedades Raras (CIBERER U-723), Madrid, Spain.
17. Unidad de ELA, Instituto de Investigación Hospital Gregorio Marañón de Madrid, SERMAS, Spain.
18. Neurology Clinic, Clinical Center of Serbia School of Medicine, University of Belgrade, Serbia.
19. Department of Health Sciences, University of Eastern Piedmont, Novara, Italy.
20. ALS Center, the Azienda Ospedaliero Universitaria Maggiore della Carità, Novara, Italy.
21. Neurology Unit, IRCCS Foundation Ca' Granda Ospedale Maggiore Policlinico, Milan, Italy.
22. Department of Brain and Behavior, University of Pavia, 27100 Pavia, Italy and General Neurology Unit, IRCCS Mondino Foundation, 27100 Pavia, Italy.
23. Genomic and post-Genomic Center, Mondino Foundation – IRCCS, 2 - 27100 Pavia.
24. Department of Neurosciences, University of Padova, Padova, Italy.
25. Department of Neurology, Brain Center Rudolf Magnus, University Medical Center Utrecht, Utrecht, the Netherlands.
26. Department of Neuroscience, Mayo Clinic, Jacksonville, Florida, USA.
27. Department of Neurology, Mayo Clinic Florida, Jacksonville, Florida 32224, USA.
28. 3rd Neurology Unit, Motor Neuron Diseases Center, Fondazione IRCCS Istituto Neurologico 'Carlo Besta', Milan, Italy.
29. Humanitas Clinical and Research Center – IRCCS (Mi) Italy and Humanitas University, Department of Biomedical Sciences, Milan, Italy.
30. Centre for Motor Neuron Disease Research, Department of Biomedical Sciences, Faculty of Medicine and Health Sciences, Macquarie University, Sydney, New South Wales, Australia.
31. Human Genetics and Cognitive Functions Unit, Institut Pasteur, Paris, France.
32. Montreal Neurological Institute, Department of Neurology and Neurosurgery, McGill University, Montreal, Quebec, Canada.
33. Academic Unit of Neurology, Trinity Biomedical Sciences Institute, Trinity College Dublin, Dublin, Republic of Ireland.
34. Faculty of Medicine, Health and Life Sciences, Queen's University Belfast, UK.
35. School of Clinical and Experimental Medicine, University of Birmingham, Birmingham, UK.
36. Sheffield Institute for Translational Neuroscience, Department of Neuroscience, University of Sheffield, Sheffield S10 2HQ, UK.
37. Department of Bioinformatics and Computational Biology, Worcester Polytechnic Institute, Worcester, MA 01609, USA.

† deceased

**The members of The American Genome Center are:**

Clifton L. Dalgard[1,2], Adelani Adeleye[1], Anthony R. Soltis[1], Camille Alba[1], Coralie Viollet[1], Dagmar Bacikova[1], Daniel N. Hupalo[1], Gauthaman Sukumar[1], Harvey B. Pollard[1,2], Matthew D. Wilkerson[1,2], Elisa McGrath Martinez[1]

1. The American Genome Center, Collaborative Health Initiative Research Program, Uniformed Services University of the Health Sciences, Bethesda, MD 20814, USA
2. Department of Anatomy, Physiology & Genetics, Uniformed Services University of the Health Sciences, Bethesda, MD 20814, USA
